# Inhibitory parvalbumin basket cell activity is selectively reduced during hippocampal sharp wave ripples in a mouse model of familial Alzheimer’s disease

**DOI:** 10.1101/2020.02.21.959676

**Authors:** Adam Caccavano, P. Lorenzo Bozzelli, Patrick A. Forcelli, Daniel T.S. Pak, Jian-Young Wu, Katherine Conant, Stefano Vicini

**Affiliations:** Interdisciplinary Program in Neuroscience, Georgetown University Medical Center, Washington, DC, 20057; Dept. of Pharmacology & Physiology, Georgetown University Medical Center, Washington, DC, 20057; Dept. of Neuroscience, Georgetown University Medical Center, Washington, DC, 20057

## Abstract

Memory disruption in mild cognitive impairment (MCI) and Alzheimer’s disease (AD) is poorly understood, particularly at early stages preceding neurodegeneration. In mouse models of AD, there are disruptions to sharp wave ripples (SWRs), hippocampal population events with a critical role in memory consolidation. However, the micro-circuitry underlying these disruptions is under-explored. We tested if a selective reduction in parvalbumin-expressing (PV) inhibitory interneuron activity underlies hyperactivity and SWR disruption. We employed the 5xFAD model of familial AD crossed with mouse lines labeling excitatory pyramidal cells (PCs) and inhibitory PV cells. We observed a 33% increase in frequency, 58% increase in amplitude, and 8% decrease in duration of SWRs in *ex vivo* slices from male and female 3-month 5xFAD mice versus littermate controls. 5xFAD mice of the same age were impaired in a hippocampal-dependent memory task. Concurrent with SWR recordings, we performed calcium imaging, cell-attached, and whole-cell recordings of PC and PV cells within the CA1 region. PCs in 5xFAD mice participated in enlarged ensembles, with superficial PCs having a higher probability of spiking during SWRs. Both deep and superficial PCs displayed an increased synaptic E/I ratio, suggesting a disinhibitory mechanism. In contrast, we observed a 46% spike rate reduction during SWRs in PV basket cells (PVBCs), while PV bistratified and axo-axonic cells were unimpaired. Excitatory synaptic drive to PVBCs was selectively reduced by 50%, resulting in decreased E/I ratio. Considering prior studies of intrinsic PV cell dysfunction in AD, these findings suggest alterations to the PC-PVBC micro-circuit also contribute to impairment.

**Significance Statement:** We demonstrate that a specific sub-type of inhibitory neuron, parvalbumin-expressing basket cells, have selectively reduced activity in a model of Alzheimer’s disease during activity critical for the consolidation of memory. These results identify a potential cellular target for therapeutic intervention to restore aberrant network activity in early amyloid pathology. While parvalbumin cells have previously been identified as a potential therapeutic target, this study for the first time recognizes that other parvalbumin neuronal sub-types, including bistratified and axo-axonic cells, are spared. These experiments are the first to record synaptic and spiking activity during sharp wave ripple events in early amyloid pathology and reveal that a selective decrease in excitatory synaptic drive to parvalbumin basket cells likely underlies reduced function.

## Introduction

Alzheimer’s disease (AD) is the leading cause of dementia, and a growing public health crisis as worldwide life expectancy increases (Mattson, 2004). AD is characterized by learning and memory deficits, the pathological accumulation of amyloid beta (Aβ) plaques and neurofibrillary tangles, and synaptic and neuronal degeneration (Serrano-Pozo et al., 2011). The cause of memory disruption in the disease is poorly understood, particularly at early ages prior to widespread neurodegeneration. The hippocampus, a region particularly important for the encoding and consolidation of spatial memory, is one of the first regions impaired in AD (Braak and Braak, 1991). Hyperactivity within the hippocampus is observed in mouse models of AD (Palop et al., 2007; Busche et al., 2008, 2012; Palop and Mucke, 2010), as well as in clinical populations, where seizures are an increasingly recognized co-morbidity of AD (Hauser et al., 1986; Amatniek et al., 2006; Palop and Mucke, 2009). While it is well appreciated that Aβ impairs the synaptic function of excitatory pyramidal cells (PCs) in later disease progression (Kamenetz et al., 2003; Shankar et al., 2008; Pozueta et al., 2013), there is growing evidence of early deficits to inhibitory GABAergic cells (Li et al., 2016), potentially explaining this shift to hyperactivity through disinhibition. In particular, several functional impairments are observed in inhibitory parvalbumin-expressing (PV) fast-spiking interneurons (Verret et al., 2012; Mahar et al., 2016; Yang et al., 2016; Hijazi et al., 2019). However, there are at least three distinct PV cell sub-types within the CA1 region of hippocampus with varying anatomical connections and function (Varga et al., 2014), and the separate impact of AD pathology on these sub-types is unknown.

PV cells play a critical role in hippocampal sharp wave ripples (SWRs) (Ellender et al., 2010; Schlingloff et al., 2014; Ognjanovski et al., 2017), spontaneous neuronal population events characterized by a low frequency sharp wave (1-30 Hz) and a high frequency ripple (120-250 Hz) (Buzsáki, 1986, 2015; Colgin, 2016). SWRs principally originate in the CA3 region and propagate to CA1 along the Schaffer collaterals, occurring in all mammalian species studied to date (Buzsáki et al., 2013). Even following decortication in brain slices, SWRs spontaneously arise in hippocampus (Buzsáki et al., 1983). SWRs have been extensively studied in large part due to their proposed role in memory consolidation (Wilson and McNaughton, 1994; Kudrimoti et al., 1999; O’Neill et al., 2008; Karlsson and Frank, 2009).

Sequences of place cells activated during spatial learning replay in temporally compressed neuronal ensembles within SWRs during rest (Nádasdy et al., 1999; Lee and Wilson, 2002). Online interruption of SWRs through both electrical (Girardeau et al., 2009; Ego-Stengel and Wilson, 2010) and optogenetic stimulation (Ven et al., 2016; Roux et al., 2017) leads to learning and memory deficits, demonstrating their critical role in memory consolidation. Notably, in several mouse models of AD, SWRs are disrupted (Gillespie et al., 2016; Iaccarino et al., 2016; Nicole et al., 2016; Xiao et al., 2017; Hollnagel et al., 2019; Jones et al., 2019; Jura et al., 2019). However, the micro-circuitry underlying these disruptions has yet to be explored in detail. In this study we employed the 5xFAD mouse model of familial Alzheimer’s disease crossed with mouse lines that selectively fluoresce in excitatory pyramidal cells (PCs) and inhibitory PV cells. We performed patch clamp recordings of deep and superficial PCs and three distinct PV cell sub-types to record the spiking activity and synaptic input during SWR events in *ex vivo* slices. Our findings support the hypothesis that a preferential reduction in synaptic input and activity of PV basket cells underlies downstream network alterations and suggest that long-term alterations to PC-PVBC micro-circuitry contribute to dysfunction in early amyloid pathology.

## Materials and Methods

### Experimental Animals

To record the activity of excitatory PCs and inhibitory PV cells in amyloid pathology, we employed a combined breeding strategy of transgenic and targeted knockin mice. Transgenic 5xFAD mice (RRID:MMRRC_034840-JAX) (Oakley et al., 2006) were back-crossed for over five generations to the C57BL/6J (RRID:IMSR_JAX:000664) background, which was common to all other strains used. To target the calcium activity of PCs under confocal microscopy, transgenic homozygous Thy1-GCaMP6f-GP5.5 (RRID:IMSR_JAX:024276) (Dana et al., 2014) were crossed with hemizygous 5xFAD mice to yield litters with both 5xFAD/+;Thy1-GCaMP6f/+ and Thy1-GCaMP6f/+ littermate controls. PV cells were identified by crossing double homozygous knockin PV^Cre^/PV^Cre^;tdTom/tdTom (RRID:IMSR_JAX:008069, RRID:IMSR_JAX:007914) (Hippenmeyer et al., 2005; Madisen et al., 2010) with hemizygous 5xFAD mice to yield litters with both 5xFAD/+;PV^Cre^/+;tdTom/+ and PV^Cre^/+;tdTom/+ littermate controls. In a subset of experiments the reporter lines were crossed, yielding quadruple transgenic cohorts of 5xFAD/+;Thy1-GCaMP6f/+;PV^Cre^/+;tdTom/+ and Thy1-GCaMP6f/+;PV^Cre^/+;tdTom/+ littermate controls. The initial intention was to use a consistent cohort of quadruple transgenic mice for all experiments, yet the breeding strategy proved inhibitive for the number of experiments, thus, patch clamp data were pooled across reporter genotype (Table 1). 5xFAD genotype was assessed at age P7 by tail biopsy via automated genotyping services (Transnetyx, Cordoba, TN, USA). For all experiments, experimenters were blind to 5xFAD genotype until after data collection and analysis were fully complete. Mice were weaned at P21 and group housed in cages with 3-5 mice separated by sex. As a model of early amyloid pathology prior to neuronal or synaptic loss (Oakley et al., 2006), two experimental cohorts were chosen (each including both males and females) at 1 month (mo) and 3 mo of age. Mice were kept on a standard 12 hr light/dark cycle, food and water were provided *ad libitum*, with all experimental procedures performed in accordance with the guidelines of the Georgetown University Animal Care and Use Committee.

**Table 1:** Mouse cohorts used in study. *Top*, Electrophysiology/imaging cohorts were distinct from behavioral cohorts. For both experiments, mice were studied at an age of 1 and 3 months (mo). *Bottom*, The 3 mo electrophysiology/imaging cohort consisted of three sub-cohorts with different reporter genes.

| Electrophysiology + Imaging (Fig. 2-8) |  |  |  |  | Behavior (Fig. 1) |  |  |  |
| --- | --- | --- | --- | --- | --- | --- | --- | --- |
|  | *Control 1mo* | *5xFAD 1mo* | ***Control 3mo*** | ***5xFAD 3mo*** | *Control 1mo* | *5xFAD 1mo* | *Control 3mo* | *5xFAD 3mo* |
| *n<sub>mice</sub>* | 10 (6f) | 10 (4f) | 27 (17f) | 29 (12f) | 10 (2f) | 10 (6f) | 15 (7f) | 12 (8f) |
| *Age (postnatal days)* | 40.2 ± 4.4 | 40.4 ± 4.6 | 95.8 ± 6.0 | 94.5 ± 5.8 | 42.0 ± 5.2 | 43.8 ± 4.2 | 91.8 ± 5.2 | 92.5 ± 5.4 |
| *n<sub>slice</sub> (range 2 - 8)* | 5.6 ± 0.8 | 4.0 ± 1.8 | 3.9 ± 1.3 | 3.2 ± 1.4 |  |  |  |  |
| *% slices with SWRs* | 93.8 ± 8.1% | 70.8 ± 22.0% | 72.4 ± 19.8% | 66.3 ± 26.1% |  |  |  |  |
|  | Thy1-GCaMP6f (Fig. 3-5) |  | PV<sup>Cre</sup>-tdTom (Fig. 6-8) |  | Thy1- GCaMP6f;PV<sup>Cre</sup> - tdTom (Fig. 4- 8) |  |  |  |
|  | *Control* | *5xFAD* | *Control* | *5xFAD* | *Control* | *5xFAD* |  |  |
| *n<sub>mice</sub>* | 10 (5f) | 11 (5f) | 13 (8f) | 11 (5f) | 4 (4f) | 7 (2f) |  |  |
| *Age (postnatal days)* | 95.0 ± 7.4 | 93.4 ± 3.2 | 96.8 ± 5.7 | 97.4 ± 6.0 | 94.8 ± 3.2 | 91.6 ± 2.1 |  |  |
| *n<sub>slice</sub> (range 2 - 8)* | 3.9 ± 1.1 | 2.8 ± 1.3 | 3.8 ± 1.5 | 4.9 ± 2.1 | 4.5 ± 1.3 | 3.6 ± 1.8 |  |  |
| *% slices with SWRs* | 75.5 ± 20.2% | 67.9 ± 26.2% | 67.4 ± 20.5% | 68.2 ± 25.3% | 80.8 ± 16.4% | 60.7 ± 30.5% |  |  |
| *n<sub>cell-attached</sub>* | 39 (PC) | 39 (PC) | 13 (PV) | 18 (PV) | 11 (PV) | 5 (PC); 14 (PV) |  |  |
| *n<sub>whole-cell</sub>* | 21 (PC) | 15 (PC) | 11 (PV) | 16 (PV) | 10 (PV) | 1 (PC); 13 (PV) |  |  |

### Amyloid Staining

Mice were anesthetized with isoflurane, transcardially perfused in iced (0° C) PBS and fixed for 48 hrs in PFA at 4° C. Brains were then given four 15 min PBS washes, and sliced horizontally at 100 μm thickness with a Vibratome Series 1000. In free floating slices, antigen retrieval was performed for 20 min in a steamer with citrate buffer containing 10 mM Na_3_C_6_H_5_O_7_, 0.05% Tween20, and pH adjusted to 6.0 with HCl. Slices were cooled to room temperature in PBS, then permeabilized for 30 min with 0.5% Triton-X in PBS, given two more PBS rinses, blocked for 2 hrs with 10% NGS and 5% BSA in PBS, and incubated overnight at 4° C with the primary monoclonal mouse antibody MOAB-2 against beta amyloid (1:500; Abcam Cat# ab126649) (Youmans et al., 2012), 0.1% Tween20, 1% NGS, 1% BSA in PBS. The following day slices were given four 15 min PBS washes and incubated in secondary containing Alexa Fluor^®^ 647 conjugated goat anti-mouse IgG (1:500; Jackson Immunoresearch Cat# 115-605-003, RRID:AB_2338902), Thioflavin-S (1:2000; Sigma-Aldrich Cat# T1892), 1% NGS in PBS for 2 hrs at room temperature. Slices were given three more 15 min PBS washes, rinsed in ddH_2_0, and mounted on slides with Vectashield® antifade mounting medium with DAPI (Vector Laboratories Cat# H-1200, RRID:AB_2336790).

### Behavioral Testing

Hippocampal-dependent learning and memory deficits were assessed by the Barnes maze (Barnes, 1979), with some modifications. The Barnes maze was conducted on a white plastic apparatus (San Diego Instruments) 0.914 m in diameter, with overhead bright illumination (286 lx) serving as the aversive stimulus. The target hole was randomly selected, with four distal visual clues present for visuospatial learning. The training phase consisted of four 180 s trials per day for 4 consecutive days. The probe trial, in which the target hole was inaccessible, was conducted on the fifth consecutive day, consisting of one 90 s trial. Mice were tracked using the ANY-maze tracking system, which was used for distance and speed measurements. The primary measures of latency and number of entries to target hole were hand-scored for increased reliability. Anxiety-like behavior was tested on the elevated plus maze (Pellow et al., 1985). The mice could explore the maze for 5 min, in which time the number of entries and fraction of time in the open arms were assessed by the ANY-maze tracking system. All tests were conducted in the light cycle, at consistent times of the day for each mouse, in an enclosed behavior room with 50 dB ambient sound and 23 lx ambient illumination. Males and females were run on the same day, but in separate groups. Cohorts were not balanced by both sex and genotype, but as much as possible testing order was prepared so that control and 5xFAD mice were alternated.

### Acute Slice Preparation

Brain slices were prepared using NMDG and HEPES-buffered artificial cerebrospinal fluid (aCSF) following a protective recovery protocol (Ting et al., 2014, 2018). Briefly, mice were anesthetized with isoflurane, transcardially perfused, dissected, and sliced in iced (0° C) NMDG-aCSF containing in mM: 92 NMDG, 2.5 KCl, 1.25 NaH_2_PO_4_, 30 NaHCO_3_, 20 HEPES, 25 Glucose, 10 Sucrose, 5 Ascorbic Acid, 2 Thiourea, 3 Sodium Pyruvate, 5 N-acetyl-L-cysteine, 10 MgSO_4_, 0.5 CaCl_2_, pH to 7.3-7.4 with HCl (300-310 Osm). All common reagents were obtained from Thermo Fisher Scientific. The brains were sliced horizontally at 500 μm thickness with a Vibratome Series 3000 to preserve hippocampal micro-circuitry and spontaneous SWRs. 3-4 slices were typically obtained per brain, which were bisected so that 6-8 hemislices in total were studied per animal. Slices spanned the dorsal-ventral axis, though were primarily medial, as only horizontal slices with intact DG, CA3, and CA1 were retained, ranging from bregma 2-4 mm. The slices were transferred together to heated (33° C) NMDG-aCSF, in which Na^+^ was gradually introduced along an increasing concentration gradient every 5 min before transferring to room temperature HEPES-aCSF containing in mM: 92 NaCl, 2.5 KCl, 1.25 NaH_2_PO_4_, 30 NaHCO_3_, 20 HEPES, 25 Glucose, 5 Ascorbic Acid, 2 Thiourea, 3 Sodium Pyruvate, 5 N-acetyl-L-cysteine, 2 MgSO_4_, 2 CaCl_2_, pH to 7.3-7.4 with NaOH (300-310 Osm). Slices recovered for 4 hours in a custom-built 150 mL incubation chamber with circulating oxygenated HEPES-aCSF.

### Slice Electrophysiology

Slices were transferred to a Siskiyou PC-H perfusion chamber with a custom-built suspended Lycra thread grid to allow perfusion below and above slice, modeled after (Hájos et al., 2009). Submerged slices were anchored with Warner Instruments slice anchors so that they were sandwiched between two grids, and perfused at a rate of 5 mL/min with heated (30° C) oxygenated aCSF containing in mM: 124 NaCl, 3.5 KCl, 1.2 NaH_2_PO_4_, 26 NaHCO_3_, 10 Glucose, 1 MgCl_2_, 2 CaCl_2_, pH 7.3-7.4 (300-310 Osm). Recordings were conducted with a Multiclamp 700B amplifier (Molecular Devices), digitized at 20 kHz and low-pass Bessel-filtered at 2 kHz with a personal computer running Clampex 11 and a DigiData 1440 (Molecular Devices). Two concurrent channels were captured: the local field potential (LFP) was recorded with 0.5-1 MΩ borosilicate pipettes filled with aCSF, paired with 3-5 MΩ borosilicate pipettes for cellular recordings. Cell-attached recordings were performed with aCSF + 5 μM Alexa Fluor^®^ 488 nm (Molecular Probes Cat# A-10440) or 594 nm (Molecular Probes Cat# A-10442) for pipette localization under confocal microscopy. The selected concentration of Alexa dye was below reported alterations to synaptic transmission (Maroteaux and Liu, 2016). Cell-attached recordings were followed with whole-cell recordings of the same cell with a new pipette filled with a Cesium internal, containing in mM: 120 CsMeSO_3_, 5 NaCl, 10 TEA•Cl, 10 HEPES, 1.1 EGTA, 4 QX314, 4 ATP•Na, 0.3 GTP•Na, pH to 7.2 with CsOH (285 Osm). For pipette localization and *post hoc* morphological reconstruction, 5 μM Alexa (either 488 or 594 nm) and 0.5% wt/vl biocytin were added to the internal solution on the day of experiment.

The LFP electrode was placed in CA1 on the border of *stratum pyramidale* (*str. pyr.*) and *oriens*, a location where both high amplitude sharp waves (SWs) and ripples are simultaneously detectable. Consistent placement of the electrode was attempted in all slices at a depth of ∼20 µm. Recordings began 10 min after LFP electrode placement to allow slice to recover. If visually detectable SWRs were not observed, the slice was logged as non-SWR producing (Table 1) and discarded. A fluorescent cell was targeted for a loose (20-40 MΩ seal resistance) cell-attached recording of 3-5 min duration. For Thy1-GCaMP6f slices, Ca^2+^ ensemble activity was recorded concurrently with a laser scanning confocal microscope system (Thor Imaging Systems Division) equipped with 488/561/642 nm lasers and green/red/far-red filters and dichroics mounted on an upright Eclipse FN1 microscope (Nikon Instruments). One thousand 512 × 512 pixel frames were captured at a sample rate of 7.5 Hz. A 40x water immersion objective was used, covering an imaging field of 350 × 350 μm, as a balance between maximizing the imaging field while providing sufficient magnification for patch clamp electrophysiology.

Following the cell-attached recording, the same cell was targeted with a new Cesium internal electrode. Upon reaching 1 GΩ seal resistance, the membrane was broken by voltage pulse and quick negative pressure. Access resistance was monitored periodically and recordings with a change >20% were discarded. Putative excitatory post-synaptic currents (EPSCs) were measured in voltage-clamp at a holding voltage of −70 mV, and putative inhibitory post-synaptic currents (IPSCs) in the same cell at 0 mV. Glutamatergic and GABAergic events were not pharmacologically isolated, as the primary goal was to correlate synaptic activity with spontaneous SWRs, which would be affected by drug administration. The reversal potentials of putative EPSCs and IPSCs matched closely with the expected glutamate and GABA reversal potentials of 0 and −70 mV. This was determined by holding the cell from +20 to −100 mV in 10 mV voltage steps and monitoring the current polarity inversion. Each voltage-clamp recording ranged from 1-3 min. Following PV cell voltage-clamp recordings, the cell was then switched to current-clamp, with current injection to offset the leak current and maintain a membrane potential of −70 mV. 35 hyperpolarizing/depolarizing steps of 5 pA increments were delivered to fill the cell with biocytin. A total duration of at least 15 min of whole-cell configuration was maintained, after which an outside-out patch was formed by slowly withdrawing the pipette. Recordings were attempted at −70 and 0 mV for all PCs, while only a subset of PV cells were clamped at 0 mV. The PV cell protocol was initially designed to minimize cell disruption and ensure proper biocytin-filling; however, the recording protocol was revised during the experiment to also record inhibitory input.

### Post Hoc Staining and Microscopy

Slices with biocytin-filled PV cells were returned to the HEPES-aCSF incubation chamber for the remainder of the day, then fixed overnight in 4% PFA, 4% glucose in PBS at 4° C. Slices then received four 15 min PBS washes, 2 hours of permeabilization with 0.5% Triton-X in PBS, 2 PBS rinses, 3 hours in Fluorescein-Avidin (1:500; Vector Laboratories Cat# A-2001, RRID:AB_2336455) in PBS, and four additional 15 min PBS washes. Free floating slices were imaged with a laser scanning confocal microscope system (Thor Imaging Systems Division). Under 20x magnification, z-stacks were obtained covering the span of visible cellular processes (40-80 µm in 1 µm steps), for both green (biocytin) and red (PV^Cre^-tdTom) channels. In a subset of 14 slices (n_slice_ = 5 Control (CT), 9 5xFAD from n_mice_ = 4 CT, 5 5xFAD), an additional round of immuno-staining was performed for Ankyrin G, which labels the axon initial segment of pyramidal cells. A separate subset of Thy1-GCaMP6f slices (n_slice_ = 2 CT, 2 5xFAD each from a separate animal) were stained for Calbindin, with differential expression between superficial and deep PCs. Antigen retrieval was performed on the 500 μm slices for 20 min in a steamer with citrate buffer. Slices were cooled to room temperature for 20 min in PBS, then blocked overnight with 10% NGS at 4° C. The following day the slices were incubated with a primary monoclonal mouse antibody against Ankyrin G (1:100; Thermo Fisher Scientific Cat# 33-8800, RRID:AB_2533145) or Calbindin D-28k (1:1000; Swant Cat# 300, RRID:AB_10000347), 0.1% Tween20, 1% NGS in PBS. After 48 hrs of primary incubation at 4° C, slices were given four 15 min PBS washes and incubated in secondary containing Alexa Fluor^®^ 647 conjugated goat anti-mouse IgG (1:500; Jackson Immunoresearch Cat# 115-605-003, RRID:AB_2338902) and 1% NGS in PBS for 3 hrs at room temperature. Slices were given four more 15 min PBS washes, rinsed in ddH_2_0, and mounted on slides between silicone isolators with Vectashield® mounting medium.

### Pre-processing of Electrophysiology Data and Event Detection

Pre-processing of files was conducted in Clampfit 11 (pClamp, Molecular Devices). Files for calcium imaging experiments were trimmed around the confocal laser trigger signal for alignment of SWR and calcium transients. Spikes were detected in cell-attached recordings with a 3 ms or 1.75 ms template search for PCs or PV cells, respectively. A threshold of 6-8 was used (Clements and Bekkers, 1997) and false negatives minimized (confirmed by manual inspection). False positives were removed by plotting peak vs anti-peak amplitude to segregate noise from true spikes. Bursts were detected with the built-in Burst Analysis, defined as three or more successive spikes, each within 60 or 40 ms (intra-burst interval) for PCs or PV cells, respectively. Spike start, peak, and end times, burst start and end times, number of events in burst, and intra-burst interval were exported from Clampfit for coincidence detection in MATLAB.

Whole-cell recordings of post-synaptic current (PSC) signals were low-pass filtered below 1000 Hz with a zero-phase Gaussian FIR filter. PSCs were detected with template searches. Multiple template categories (3-4) of varying duration (ranging from 3-20 ms) were used to improve detection for overlapping PSCs often seen around SWRs. Shorter duration templates used increasingly higher thresholds, between 5-8, to minimize false negatives. Parameters were kept constant for each cell type. PSC results including start, peak, and end times, amplitude, rise tau and decay tau were exported from Clampfit and processed in Microsoft Excel to remove duplicate events with identical peak times. Limits were set on rise and decay tau to remove false positives due to noise. Accepted EPSCs fell within a range of 0.05-5 ms rise tau and 1-50 ms decay tau, while IPSCs fell within a range of 0.1-10 ms rise tau and 3-100 ms decay tau.

### Local Field Analysis and Sharp Wave Ripple Detection

All analysis for coincidence detection between the LFP and cellular events was conducted with custom-built MATLAB functions. Raw traces were imported using the abfload protocol (https://github.com/fcollman/abfload). All applied filters were finite impulse response (FIR) Gaussian filter with constant and corrected phase delays. A wide band-pass (1-1000 Hz) was first applied to remove both low frequency DC drift and high frequency instrument noise. The detection of SWR events was based upon prior approaches (Siapas and Wilson, 1998; Csicsvari et al., 1999; Eschenko et al., 2008), with refinements to minimize false positives and permit additional analyses. The LFP was filtered in both the sharp wave (SW: 1-30 Hz) and ripple (120-220 Hz) ranges, and the root mean square (RMS) was computed every 5 ms in a 10 ms sliding window. The threshold for peak detection was set to 4 standard deviation (SD) above the baseline (lower 0.95 quantile) RMS mean. Event start and end times were set at 2 SD crossings. SWR events were defined as the intersection of concurrent SW and ripple events. The duration of SWR events was determined from the union of concurrent SW and ripple events. The peak of the SWR event was defined as the peak of the SW-RMS signal, and the amplitude as the difference between the peak and baseline values of the SW signal. The power of the SW and ripple were determined by the bandpower MATLAB function of the relevant filtered signal, which computes an approximation of the integral of the power spectral density between the start and end times of the SWR event. Additional filters were applied in the slow gamma (20-50 Hz) and fast/pathological ripple (250-500 Hz) ranges, and the power computed on a SWR-event basis, as the power of these SWR-nested oscillations has been implicated in memory performance (Carr et al., 2012) and epileptogenesis (Foffani et al., 2007), respectively.

To visualize the spectral components of the LFP as a function of time, spectral analysis was performed for the duration of the recording as well as per SWR event via a Short-Time Fourier Transform (STFT) between 1-500 Hz. To better observe deviations from baseline power, the Z-score for each 1 Hz frequency band was calculated. A Fast-Fourier Transform (FFT) was also computed for a 200 ms window centered around each SWR peak and averaged across all events as an additional visualization of spectral power. The determination of the phase of slow gamma and ripple oscillations during SWRs was based on the analysis of (Varga et al., 2012). Within a 200 ms window centered around the SWR peak, the extrema of the filtered signal of interest were identified, and a piece-wise linear function fit with values from 0° to 180° between a maximum and minimum, 180° to 360° between a minimum and maximum, and then resetting to 0°. The number of cycles within the duration of the SWR event was recorded, from which the peak frequency was calculated as *n*_*cycle*_*/SWR*_*duration*_.

### Pre-processing of Ca^2+^ Imaging Data

Raw time series were converted to the change in fluorescence normalized to baseline (*ΔF/F*) with custom-built ImageJ (FIJI) macros, in which batches of raw TIF images were imported with the Bio-Formats plugin (Open Microscopy Environment) and saved as TIF stacks. The TIF stacks were corrected for photo-bleaching via two iterations of the built-in Correct-Bleach plugin. Photo-bleaching was modeled as a sum of two exponentials, a fast 5 s decay and a slower decay over the duration of the recording (133 s). To assist in region-of-interest (ROI) placement, a semi-automated algorithm was applied to highlight regions of highest fluorescence change. For each TIF stack, the squared coefficient of variation (SCV) image was calculated, defined as the variance divided by the average squared for each pixel, or equivalently:

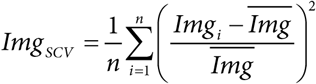

Circular ROIs were manually drawn over all identified cells in the SCV image and confirmed with the time series to encompass active cells. Notably, in this way only cells with variable fluorescence were identified, and static highly fluorescent Ca^2+^-loaded cells were excluded. The *ΔF/F* for each cell was calculated by first subtracting the background fluorescence *F*_*b*_, defined as the lowest intensity pixel across the entire time series. *F*_*0*_ for each ROI was defined as the average of the ten images with the lowest intensity. The *ΔF/F* was thus calculated as:

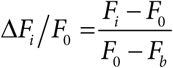

These values were exported from ImageJ, along with a separately calculated timing file for each image, computed from a threshold search in Clampfit of the confocal laser trigger channel.

### Calcium Transient Detection and SWR Coincidence Analysis

Automatic detection of calcium transients was performed by first correcting for slow changes in fluorescence, either from remaining photo-bleaching or gradual drift of the imaging plane. A smoothed moving average, calculated with robust locally-weighted regression (Cleveland, 1979) with a window 25% of the file duration (33 s), was subtracted from each *ΔF/F* trace. The baseline-corrected *ΔF/F* traces were interpolated from the raw sampling rate of 7.5 Hz to 2 kHz, and the LFP downsampled from 20 kHz to 2 kHz for coincidence detection. Automatic threshold detection for each cell was set at 4 SD above baseline, with the start and end times for each event set at 2 SD. The baseline of each cell was determined by an iterative algorithm of gaussian fitting to the histogram of all data points, which for most cells was a skewed one-tailed distribution with a large baseline peak at zero and a long positive tail representing transient events. For uncommon cells without a skewed distribution (kurtosis < 0), a single gaussian fit was applied to the entire histogram, providing an estimate of the baseline mean and SD. However, for most cells, two iterations were employed, with the first a double-gaussian fit to the entire histogram. The higher amplitude gaussian SD was then used to constrain the upper limit on a second iteration single-gaussian refinement fit of only the baseline. Special logic was necessary in rare situations. For excessively skewed distributions (high *ΔF/F*, kurtosis > 5) the double-gaussian fit was constrained to the lower 0.95 quantile of data. For extremely active cells, because of the slow decay kinetics of GCaMP, the signal peak rivaled or exceeded the baseline peak in amplitude. For these cells, the more negative rather than the higher amplitude peak was used as an estimate of the baseline. In the situation where two peaks were not clearly differentiated (peak separation < 0.1 *ΔF/F*), the gaussian means and SDs were averaged together to arrive at an estimate for constraining the second iteration.

Once the baseline mean, SD, and thresholds were determined for each cell, events were detected on a cell-by-cell basis, and characteristics calculated including start, peak, and end times, IEI, duration, amplitude, and frequency. The interpolated calcium traces were trimmed and aligned with the down-sampled LFP trace. Each calcium transient was classified as SWR-coincident if there was any overlap between the start and end of the calcium transient and the start and end of the SWR, otherwise it was classified as spontaneous. Cellular participation during SWRs was assessed by constructing a simplified event matrix with SWRs in one dimension, and cells in the other, with a zero or one if the cell reached threshold during the SWR. For fields with five or more active cells, the ensemble diversity was assessed by calculating the pairwise Jaccard Similarity index between SWR events, ranging from a value of zero if two SWR events had no cells in common, to a value of one if all cellular participants were identical, modeled after the analysis of (Miyawaki et al., 2014). The cell-cell pairwise index was also calculated, ranging from zero if two cells never participated in the same SWR events, to one if they participated in precisely the same SWRs. The cumulative distribution functions were determined for each recording by considering all off-diagonal values in one half of the symmetric similarity matrices.

### Spike and SWR Coincidence Analysis

Each spike/burst was classified as SWR-coincident if there was any overlap between the start and end of the spike/burst and the start and end of the SWR, otherwise it was classified as spontaneous. To examine spike rate in more detail around SWR events, the peri-SWR spike probability was calculated by sorting all spikes that occurred within a 200 ms window centered around each SWR peak into 2 ms bins and normalizing to all SWR events. Spike-phase coupling was determined by identifying the previously calculated gamma and ripple phases at the spike peak time. Spike phase times were only considered for further analysis if trough-peak amplitude exceeded 4 SD of the gamma or ripple signal.

### Post-Synaptic Current and SWR Coincidence Analysis

Each EPSC/IPSC was classified as SWR-coincident (swrEPSC/swrIPSC) if there was any overlap between the start and end of the event and a 100 ms window centered around each SWR peak, otherwise it was classified as spontaneous (sEPSC/sIPSCs). The more conservative window to classify spontaneous events (compared to the calcium and spike analysis) better captured the buildup of EPSCs/IPSCs preceding SWRs (Schlingloff et al., 2014). As events overlap and complicate detection during SWRs, for quantification the current was integrated in a 100 ms window centered at the SWR peak to determine the total charge (swrEPSQ/swrIPSQ). To examine the distribution of sEPSCs/sIPSCs, the cumulative distribution function of all events was computed for each cell and averaged across all cells in a group. For a view of the temporal progression of synaptic input during SWRs, the charge was also calculated by integrating a sliding 100 ms window every 2 ms in a 200 ms window centered around the SWR peak.

### Experimental Design and Statistical Analyses

Most experiments in this study have been presented in a case-control experimental design, in which data from 5xFAD mice are compared to littermate controls. With behavioral learning on the Barnes Maze we have employed a longitudinal repeated measures (RM) design. In analyzing the impact of factors on LFP activity in a large sample of *ex vivo* slices, we have taken a factorial design. All data were tested for normality and lognormality via Shapiro-Wilk tests. If all groups were normally distributed, they were analyzed with parametric tests (unpaired t-test, n-way ANOVA, n-way RM ANOVA) and have been displayed on bar plots with error bars representing the mean ± SEM. If any group was not normally distributed, the data have been presented on box-whisker or violin plots, with lines indicating median and quartiles, and full error bars representing range. If all groups were lognormally distributed, the data were log transformed, the results of which were analyzed with parametric tests, with the log-means compared. For clarity, the original non-transformed data have been displayed in plots. If any group was neither normal nor lognormal, non-parametric statistical tests of rank were employed. Circular data (phase angles) were analyzed with a similar approach. Data were tested for non-uniformity via a Rayleigh test to determine if they could be sampled from a von Mises (circular normal) distribution. If the Rayleigh test reached significance for all groups, means were compared with the Watson-Williams test, otherwise a circular median test was performed.

The particular statistical tests used are listed in the Results. Any values cited in the text are mean ± SEM. All *post hoc* multiple comparisons used the Šidák correction (ŠC). Within each plot all individual data points are presented. No data were excluded based on their values, but only for experimental reasons (e.g. no SWRs present, excessive slice movement, unstable patch clamp recording). The *n* is indicated in the text and figure legends, and differs between experiment, either *n*_*mice*_, *n*_*slice*_, or *n*_*cell*_. Raw p-values are displayed in plots; if less than a significance level of 0.05 they are bold. Data originating from male or female mice are presented as closed or open circles, respectively. Sex differences were examined for some endpoints, but in general the experiments were insufficiently powered to determine sex differences. A power analysis of the principal experiments was performed based on preliminary data, guiding the number of animals/cells chosen.

Graphpad Prism 8 was used for all 1 and 2-sample statistical tests. Microsoft Excel and MATLAB 2019 were used for some simple calculations of mean, SD, SEM, ratio, and error propagation. n-way ANOVAs were performed in the Statistics and Machine Learning Toolbox of MATLAB 2019. Non-parametric factorial data were aligned and ranked with ARTool (Wobbrock et al., 2011), before running ANOVAs. Circular statistics were run in the Circular Statistics Toolbox for MATLAB (Berens, 2009).

### Code Accessibility

All code is open-source and available in public repositories, including versions under active development (Github) as well as archival copies used for this manuscript (Zenodo). MATLAB functions are available at https://github.com/acaccavano/SWR-Analysis (archival copy: DOI: 10.5281/zenodo. 3625236). ImageJ (FIJI) macros are available at https://github.com/acaccavano/deltaFoF (archival copy: DOI: 10.5281/zenodo.3625130).

## Results

### Three month 5xFAD mice deposit amyloid and display minor impairment in spatial memory

The 5xFAD model of familial Alzheimer’s disease is an aggressive though useful model of amyloid pathology, as it exhibits heavy amyloid accumulation in hippocampus and associated cortices, and is accompanied with memory impairment (Oakley et al., 2006). 5xFAD mice are unimpaired in performance on the T-maze at 2 months (mo) but become impaired by 4-5 mo when amyloid burden is greater (Oakley et al., 2006). Interestingly, at 3 mo there is no evidence for neuronal or synaptic degeneration, while several synaptic markers begin to decline at 4 mo and are significantly reduced from controls by 9 mo. We observed intracellular amyloid accumulation in the subiculum and CA1 region of hippocampus in 1 mo 5xFAD mice without the presence of extracellular plaques (n_mice_ = 2 Control (CT), 2 5xFAD), while in 3 mo 5xFAD mice we observed multiple extracellular plaques in subiculum, with sparse plaques in CA1 (n_mice_ = 2 CT, 2 5xFAD, Fig. 1A). Prior work indicates that amyloid burden continues to increase throughout the lifespan of 5xFAD mice (Oakley et al., 2006; Youmans et al., 2012).

**Figure 1:**
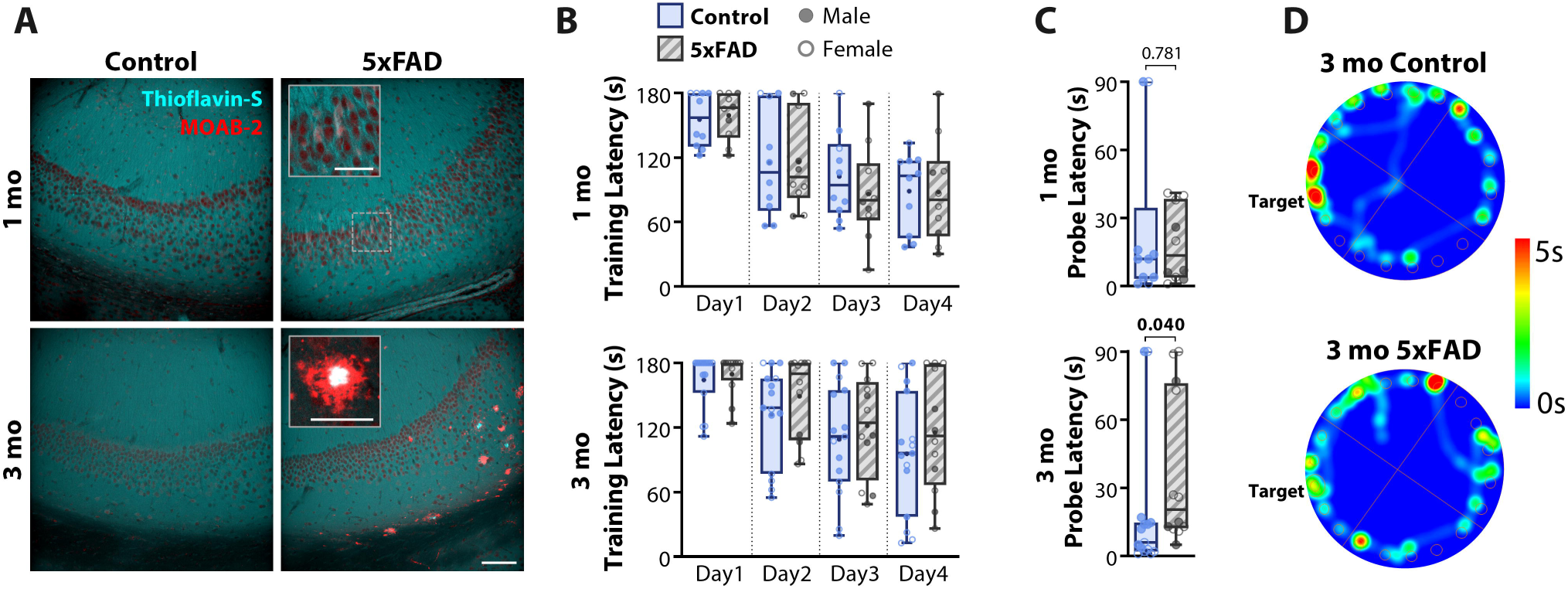
Three month (mo) 5xFAD mice deposit amyloid and display minor impairment in spatial memory. ***A***, Representative IHC in the CA1 – subiculum region in 1 mo (*top*) and 3 mo (*bottom*) for control (CT, *left*) and 5xFAD (*right*) mice. Red: MOAB-2, labeling both intraneuronal and extracellular Aβ, including unaggregated, oligomeric, and fibrillar Aβ_42_ and unaggregated Aβ_40_. Cyan: Thioflavin-S, labeling fibrils in extracellular plaques. Staining was repeated in 2 mice of each genotype and age. Scale bar = 100 μm. Inset scale bars = 50 μm. ***B***, Cohorts of 1 mo (n_mice_ = 10 CT, 10 5xFAD, *top*) and 3 mo (n_mice_ = 15 CT, 12 5xFAD, *bottom*) mice, were trained on the Barnes Maze, consisting of four days of training, each with four 180 s trials. The latency to find the target escape hole was averaged over the four trials. ***C***, Latency to find target on the final 90 s probe trial on the fifth day for 1 mo (*top*) and 3 mo (*bottom*) cohorts. ***D***, Representative heat map for 3 mo animals on the final probe trial. Warmer colors represent longer duration. Control mice spent more time at the target and surrounding region than 5xFAD mice. For all plots, individual data points represent an animal. Closed circles represent males, open circles females. Box-whisker plots represent non-normal data as Median and IQRs. p-values indicated above brackets.

In separate 1 mo (n_mice_ = 10 CT, 10 5xFAD) and 3 mo cohorts (n_mice_ = 15 CT, 12 5xFAD, Table 1), we examined the performance of 5xFAD mice and littermate controls on the Barnes Maze. Both age cohorts learned the task, as seen by decreased latency to find escape hole over progressive days of training, and a significant effect of training day (1 mo: F_(3,54)_ = 23.9, p = 5.7 × 10^−10^; 3 mo: F_(3,75)_ = 31.6, p = 2.7 × 10^−13^; 2-way Repeated Measures (RM) ANOVA, align-rank transformed (ART); Fig. 1B). There were no differences observed in training between genotype for either the 1 mo cohort (F_(1,18)_ = 0.096, p = 0.760) or the 3 mo cohort (F_(1,25)_ = 1.086, p = 0.307), nor were there interactions of genotype × training day (1 mo: F_(3,54)_ = 0.496, p = 0.687; 3 mo: F_(3,75)_ = 0.983, p = 0.405). On the probe day, 1 mo 5xFAD mice had similar latencies to controls (U = 46, p = 0.781; Mann-Whitney; Fig. 1C), while 3 mo 5xFAD mice had a longer latency to find the escape hole from 18.4 ± 7.6 s to 37.4 ± 9.8 s (U = 48, p = 0.040; Mann-Whitney; Fig. 1C-D). The number of entries to the area of the escape hole, another commonly reported endpoint, was not significantly different for either cohort (1 mo: t_(18)_ = 0.735, p = 0.472; 3 mo: t_(25)_ = 0.691, p = 0.496; unpaired t-tests). 5xFAD mice displayed no obvious motor impairments, as the total distance traveled did not differ between genotype at either age (1 mo: 3.98 ± 0.91 m (CT), 3.77 ± 0.49 m (5xFAD), t_(18)_ = 0.203, p = 0.842; 3 mo: 3.98 ± 0.49 m (CT), 3.40 ± 0.49 m (5xFAD), t_(25)_ = 0.816, p = 0.422; unpaired t-tests), neither did the mean speed (1 mo: 4.4 ± 1.0 cm/s (CT), 4.2 ± 0.5 cm/s (5xFAD), t_(18)_ = 0.186, p = 0.856; 3 mo: 4.4 ± 0.6 cm/s (CT), 3.8 ± 0.5 cm/s (5xFAD), t_(25)_ = 0.819, p = 0.420; unpaired t-tests).

To test if this observed memory impairment could be attributed to altered anxiety-like behavior, the cohorts were also tested on the Elevated Plus Maze. Neither age cohort showed a significant difference in fraction of time spent in open arms (1 mo: 15.4 ± 2.2% (CT), 15.1 ± 0.9% (5xFAD), t_(18)_ = 0.129, p = 0.900; 3 mo: 10.7 ± 1.0 (CT), 9.3 ± 1.5 (5xFAD), t_(25)_ = 0.848, p = 0.404; unpaired t-tests), nor in number of open arm entries (1 mo: 7.9 ± 0.6 (CT), 7.4 ± 0.6 (5xFAD), t_(18)_ = 0.570, p = 0.576; 3 mo: 9.2 ± 0.7 (CT), 7.5 ± 0.6 (5xFAD), t_(25)_ = 1.863, p = 0.074; unpaired t-tests). Therefore, we concluded that 3 mo 5xFAD mice on our genetic background had a mild spatial memory impairment. As memory impairment has been widely reported in 5xFAD mice at later ages (Oakley et al., 2006; Ohno, 2009; Tohda et al., 2012), no further memory tasks were performed.

### Sharp wave ripples are increased in 3 month 5xFAD mice

As spatial memory relies heavily on the activity of the hippocampus, and sharp wave ripples (SWRs) are critical for the consolidation of new memories (Buzsáki, 1986; Wilson and McNaughton, 1994), we next recorded spontaneous SWRs in hippocampal slices from control and 5xFAD mice (Fig. 2A-B). SWRs were recorded in the CA1 region in multiple slices for each animal and averaged in both a 1 mo cohort (n_mice_ = 10 CT, 10 5xFAD), and a 3 mo cohort (n_mice_ = 27 CT, 29 5xFAD, Table 1). While the electrode placement was kept as consistent as possible across recordings (see Methods), slight deviations in placement can result in large variance in the LFP. We attempted to control for this by recording from many slices (1 mo: n_slice_ = 51 CT, 45 5xFAD; 3 mo: n_slice_ = 101 CT, 88 5xFAD) and then averaging across slices for each animal. In the 1 mo cohort, the SWR event frequency did not differ (t_(18)_ = 0.946, p = 0.357; unpaired t-test; Fig. 2C), nor were there changes for any other LFP endpoints (data not shown). However, in the 3 mo cohort, SWR event frequency was increased in 5xFAD mice versus controls from 0.94 ± 0.07 Hz to 1.25 ± 0.08 Hz (t_(54)_ = 2.89, p = 0.006; unpaired t-test; Fig. 2C). We next verified if there was an altered percentage of slices exhibiting SWRs in 5xFAD mice (Table 1), as this could underlie observed differences in event frequency. While a 2-way ANOVA revealed a significant effect of 5xFAD genotype (F_(1,72)_ = 6.51, p = 0.013) and age (F_(1,72)_ = 5.21, p = 0.0254) on percentage of slices with SWRs, the only significant difference observed when corrected for multiple comparisons was between control slices at 1 mo and 3 mo (p = 0.039, Šidák correction (ŠC)), while 3 mo slices did not differ between genotype (p = 0.759, ŠC). Therefore, the observed differences in 3 mo mice are not likely due to altered viability of the slices.

**Figure 2:**
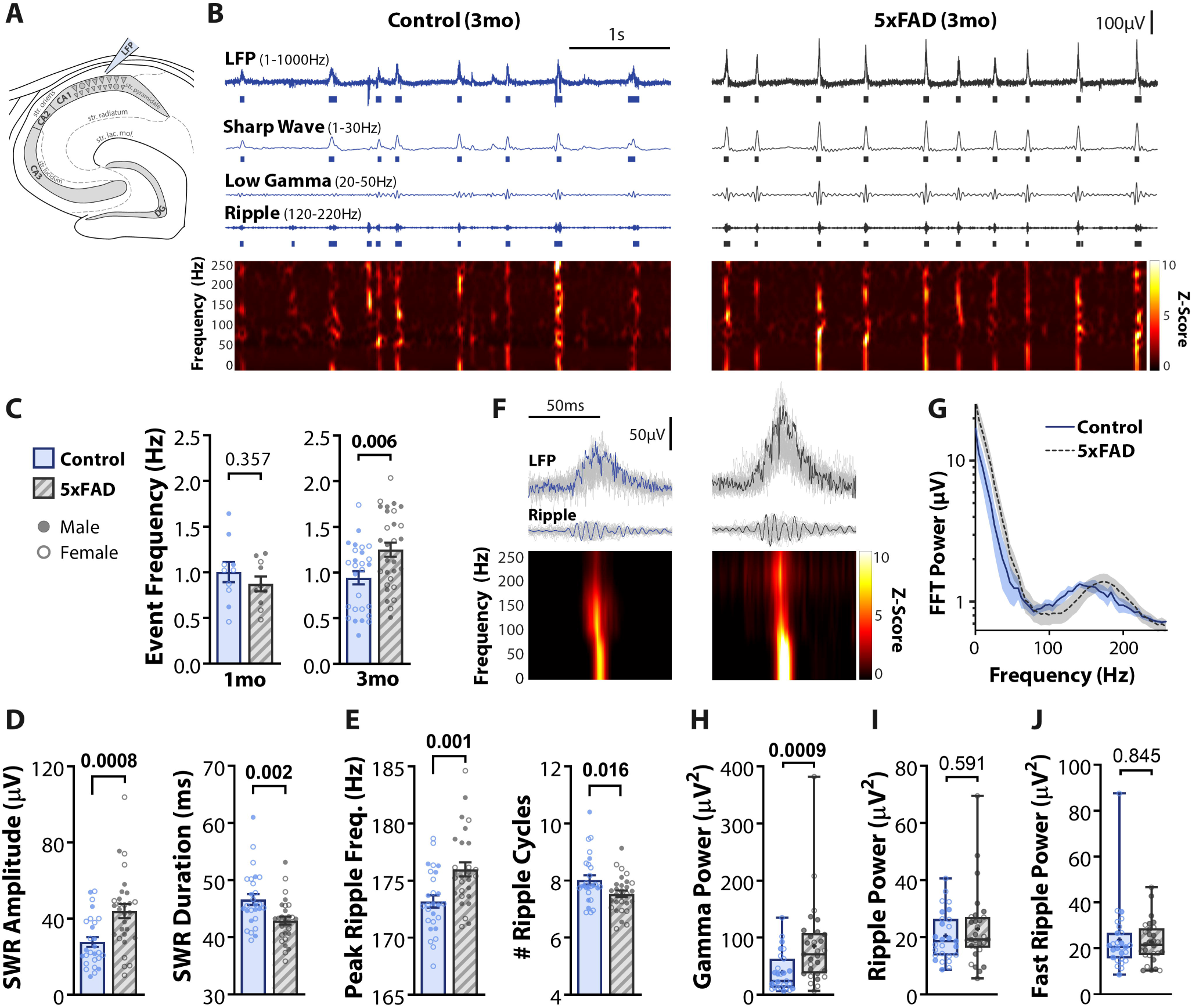
Sharp wave ripples are increased in 3 month 5xFAD mice. ***A***, SWRs were recorded in acute horizontal slices with the LFP electrode located in CA1. ***B***, Example traces of 3 mo control (*left*) and 5xFAD (*right*) slices. *1*^*st*^ *trace*, LFP filtered between 1-1000 Hz, raster below indicates detected SWR events as overlap of sharp wave (SW) and ripple events. *2*^*nd*^ *trace*, SW, filtered between 1-30 Hz, raster below indicates detected SW events. *3*^*rd*^ *trace*, low gamma, filtered between 20-50 Hz. *4*^*th*^ *trace*, ripple, filtered between 120-220 Hz, raster below indicates detected ripple events. *Bottom*, z-scored time-frequency spectrogram from 1-250 Hz. ***C***, Summary plot of SWR event frequency for 5xFAD and littermate controls for both 1 mo (n_mice_ = 10 CT, 10 5xFAD; *left*) and 3 mo cohorts (n_mice_ = 27 CT, 29 5xFAD; *right*). ***D***, SWR event amplitude and duration. ***E***, Peak ripple frequency and the number of ripple cycles within SWR duration. ***F***, Example SWR events for traces displayed in ***B***, with average z-scored time-frequency spectrogram of all events below. ***G***, Average Fast-Fourier Transform (FFT) of 200 ms window around all SWR events from characteristic subset of n_slice_ = 7 CT, 7 5xFAD from n_mice_ = 4 CT, 5 5xFAD. Shaded region represents SEM. ***H-J***, SWR-locked oscillation power in the low gamma (20-50 Hz, ***H***), ripple (120-220 Hz, ***I***), and fast ripple (250-500 Hz, ***J***) ranges. Note that in ***H***, one value was identified as an outlier by the ROUT method but was retained in analysis, as the data point appeared valid, and even with removal did not alter the observed increase (U = 193, p = 0.0015; Mann-Whitney). For all plots, individual data points represent the average of all slices recorded from an animal (n_slice_/animal in Table 1). Closed circles represent males, open circles females. Bar plots indicate normal data with Mean ± SEM. Box-whisker plots represent non-normal data with Median and IQRs. p-values indicated above brackets.

In addition to an increased event frequency, SWR amplitude was increased by 58 ± 17% in 3 mo 5xFAD mice (t_(48.9)_ = 3.59, p = 0.0008; Welch’s t-test) and surprisingly were 3.7 ± 1.2 ms shorter in duration (t_(54)_ = 3.18, p = 0.0025; unpaired t-test; Fig. 2D). This decrease in duration was likely attributable to both increased peak ripple frequency (t_(54)_ = 3.422 p = 0.0012; unpaired t-test) and decreased number of complete ripple cycles (t_(54)_ = 3.48, p = 0.016; unpaired t-test; Fig. 2E), and is of interest as longer duration SWRs have been demonstrated to improve memory (Fernández-Ruiz et al., 2019). Spectral features of SWRs also differed between genotype (Fig. 2F-J). Low-gamma (20-50 Hz) nested within SWRs, speculated to play a role in coordinating CA3 and CA1 memory replay (Carr et al., 2012), was increased in 5xFAD mice (U = 193, p = 0.0009, Mann-Whitney; Fig. 2H). While the ripple peak frequency was shifted (Fig. 2E,G), the total power within the ripple frequency range from 120-220 Hz was unchanged (U = 358, p = 0.591; Mann-Whitney; Fig. 2I). The power of fast or pathological ripples (250-500 Hz) did not differ between 5xFAD and control slices (U = 379, p = 0.845; Mann-Whitney; Fig. 2J), suggesting the increased activity in 5xFAD slices was distinct from epileptiform activity (Foffani et al., 2007).

Spontaneous SWRs recordings were repeated in three separate 3 mo sub-cohorts with different reporter mouse lines for subsequent patch-clamp and Ca^2+^ imaging experiments (Table 1). SWR frequency increased in 5xFAD mice in both larger cohorts: PV^Cre^/+;tdTom/+ mice (135 ± 18%, n_mice_ = 13 CT, 11 5xFAD) and Thy1-GCaMP6f mice (156 ± 21%, n_mice_ = 10 CT, 11 5xFAD). In the third smaller Thy1-GCaMP6f;PV^Cre^/+;tdTom/+ cohort, there was no significant increase observed (93 ± 18%, n_mice_ = 4 CT, 7 5xFAD). A 2-way ANOVA for 5xFAD genotype and reporter line revealed only a significant effect of 5xFAD genotype (F_(1,50)_ = 5.41, p = 0.024) and not reporter line (F_(2,50)_ = 1.50, p = 0.233) nor interaction term (F_(2,50)_ = 2.06, p = 0.138). The data for the three sub-cohorts were therefore pooled into the results presented in (Fig. 2), and in subsequent patch clamp experiments, with the lack of phenotype in the quadruple transgenic cohort likely attributable to smaller sampling. To test if sex, brain hemisphere, and dorsal-ventral slice position had an effect, an additional 4-way ANOVA of all slices was performed. A small effect of 5xFAD genotype was found (n_slice_ = 101 CT, 88 5xFAD; F_(1,178)_ = 10.4, p = 0.0015), and a small effect of dorsal-ventral position (F_(1,178)_ = 10.46, p = 0.0015), with ventral slices (bregma 3-4 mm) 126 ± 12% the frequency of more dorsal slices (bregma 2-3 mm) for both control and 5xFAD slices. Correcting for multiple comparisons, within each genotype the difference between dorsal and ventral slices did not reach significance (D vs V: p = 0.078 (CT), p = 0.235 (5xFAD), ŠC). No significant effects were observed for sex (F_(1,178)_ = 0.728, p = 0.395), brain hemisphere (F_(1,178)_ = 0.873, p = 0.351), nor any interaction terms.

### Altered ensembles of pyramidal cells are recruited in 5xFAD slices

The replay of pyramidal cell (PC) ensembles during SWRs is critical for the consolidation of spatial memory (Wilson and McNaughton, 1994; Skaggs and McNaughton, 1996; Lee and Wilson, 2002; O’Neill et al., 2008). Given the observed alterations to SWRs in 3 mo 5xFAD mice, we next sought to determine if ensembles of PCs were altered via calcium imaging (Fig. 3A). PCs differentiate into superficial (sPCs, closer to *str. radiatum*) and deep (dPCs, closer to *str. oriens*) cells, with different function, connectivity, and molecular profiles (Lee et al., 2014; Valero et al., 2015). We distinguished sPCs and dPCs via *post hoc* staining of imaged slices for calbindin (CB), which is more highly expressed in sPCs. While dorsal hippocampus exhibits a clear delineation of CB+ sPCs and CB-dPCs (Lee et al., 2014), we observed a CB bilayer in our slices (Fig. 3B), as previously reported in more ventral hippocampus (Baimbridge and Miller, 1982; Slomianka et al., 2011). In n_slice_ = 4 from n_mice_ = 2 CT, 2 5xFAD, we counted the total number of GCaMP (GC)+ CB-(539), GC-CB+ (292) and GC+ CB+ (146) cells, and for each calculated the distance from the border of *str. radiatum* and *pyramidale* (Fig. 3C). At 30 µm there was a switch from majority CB+ to GC+ cells, with a non-trivial fraction of double-labeled GC+ CB+ cells: 17.9% from 0 – 30 µm and 15.7% from 30 – 90 µm. At depths greater than 90 µm there was lower co-expression of 7.9%. All subsequent experiments were performed on GC+ cells with a cutoff of 30 µm between putative sPCs and dPCs.

**Figure 3:**
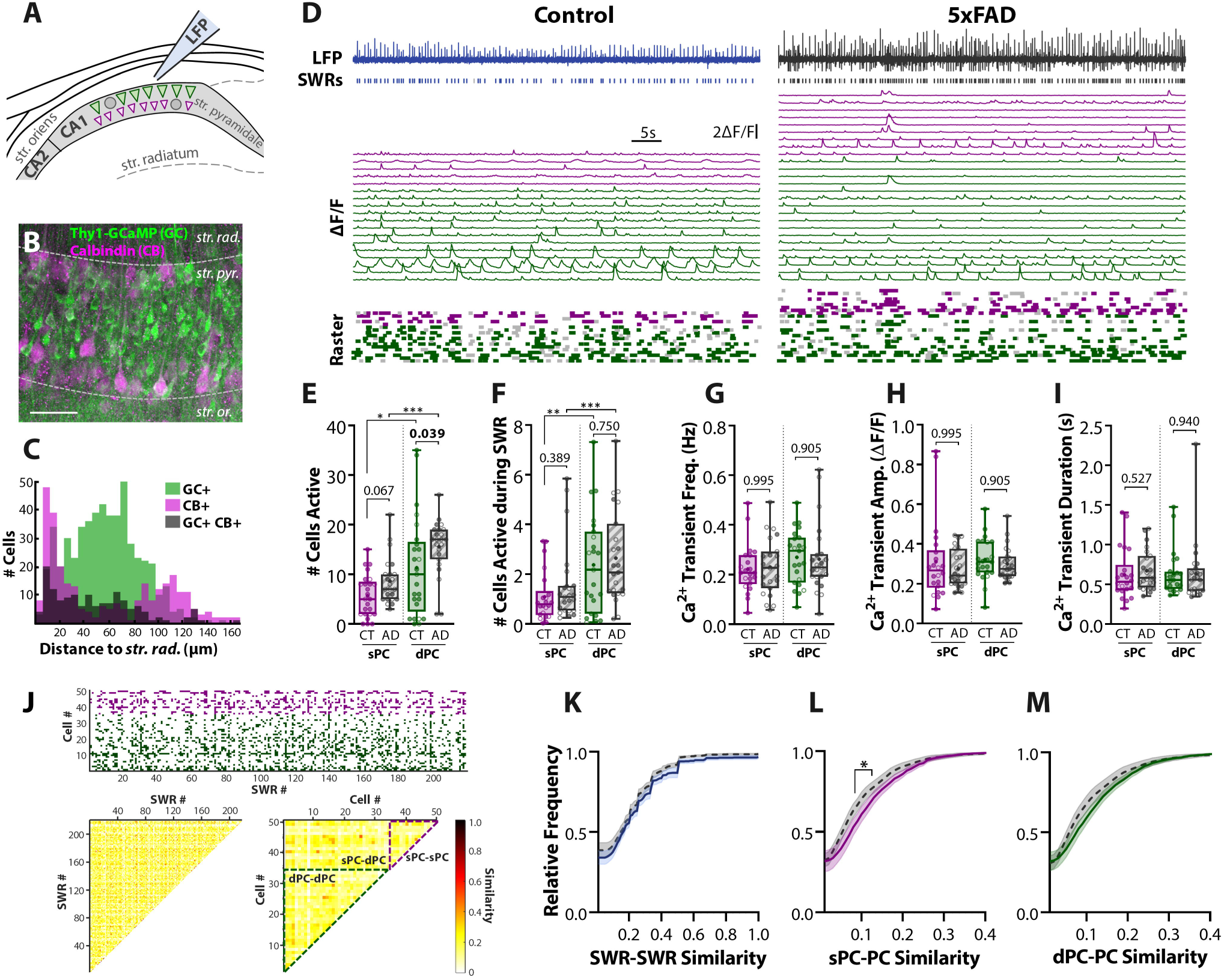
Altered ensembles of pyramidal cells are recruited in 5xFAD slices. ***A***, Confocal imaging of PC ensembles were recorded concurrently with SWRs. Scale bars = 50 μm. ***B***, *Post hoc* IHC for Calbindin (CB) of imaged Thy1-GCaMP6f (GC) slices. Dashed grey lines approximately distinguish layers of hippocampus, *str. rad.* = *stratum radiatum, str. pyr.* = *stratum pyramidale, str. or.* = *stratum oriens*. Scale bar = 50 μm. ***C***, Histogram of the distance from the center of the cell body to the *str. pyr./rad.* border for 539 GC+ CB-, 292 GC-CB+, and 146 GC+ CB+ cells pooled from n_slice_ = 4 from n_mice_ = 2 CT, 2 5xFAD. ***D***, LFP and ΔF/F for identified cells in slices from 3 mo control (*left*) and 5xFAD (*right)* mice. *Top trace*, LFP trace with identified SWR events in raster below. *Middle traces*, individual ΔF/F for each cell, superficial PCs (sPC) in magenta, and deep PCs (dPCs) in green. *Bottom raster*, identified Ca^2+^ events above threshold. Dark colored event indicate concurrence with SWR, light gray indicates spontaneous event. ***E***, The total number of active sPCs and dPCs for CT (n_slice_ = 25) and 5xFAD (n_slice_ = 23) slices. ***F***, The number of sPCs/dPCs active during SWRs. ***G***, Frequency, ***H***, Amplitude, and ***I***, Duration of Ca^2+^ transient events averaged across sPCs and dPCs for each slice. ***J***, *top*, Event matrix displaying for one example recording, the SWR events on the x-axis, and cells on the y-axis (magenta = sPC, green = dPC). *Bottom left*, pairwise Jaccard similarity between SWR events (columns in event matrix). Similarity matrix is symmetric and the diagonal = 1 by definition – these have not been plotted for clarity. *Bottom right*, pairwise Jaccard similarity between cells (rows in event matrix). Dashed lines indicate borders between sPC-sPC, sPC-dPC, and dPC-dPC comparisons. Quantification of cell similarity was performed by grouping all sPC-PC comparisons (sPC-sPC & sPC-dPC), and all dPC-PC comparisons (dPC-dPC & sPC-dPC). Cumulative distribution functions were calculated from all pairwise comparisons for ***K***, SWR-SWR similarity, ***L***, sPC-PC similarity, and ***M***, dPC-PC similar. ***K-M*** all showed significant genotype differences via 2-way ANOVAs; asterisks indicate regions surviving multiple comparisons. * p < 0.05, ** p < 0.01, *** p < 0.001. Box-whisker plots represent non-normal data with Median and IQRs. Individual data points represent a slice. Closed circles represent slices from males, open circles females. p-values of pairwise comparisons indicated above brackets.

The ensemble activity of PCs was recorded under confocal microscopy in slices from 5xFAD/+;Thy1-GCaMP6f mice and Thy1-GCaMP6f littermate controls (Fig. 3D). Active cells were detected with a semi-automated algorithm and both spontaneous and SWR-coincident Ca^2+^ transient events were detected. Within 5xFAD slices, the total number of active PCs detected per imaging field was increased from 17.0 ± 2.6 to 24.0 ± 1.6 PCs in n_slice_ = 25 CT, 23 5xFAD (t_(31.2)_ = 3.091, p = 0.0042; Welch’s t-test, log-transformed (LT)), with a greater number of PCs active during SWRs (CT: 3.6 ± 0.6, 5xFAD: 4.9 ± 0.8 cells, t_(40.1)_ = 2.063, p = 0.046; Welch’s t-test, LT). When delineated by PC sub-type, there was a significant effect of genotype on the number of active cells (F_(1_,_92)_ = 11.8, p = 0.00088; 2-way ANOVA, ART; Fig. 3E), as well as an effect of cell type (F_(1,92)_ = 20.6, p = 1.7 × 10^−5^). The number of sPCs was increased in 5xFAD slices from 5.7 ± 0.8 to 8.7 ± 1.0 cells (p = 0.067, ŠC) and dPCs from 11.3 ± 2.0 to 15.6 ± 1.2 cells (p = 0.039, ŠC), with significantly more dPCs than sPCs for both genotypes (CT: p = 0.014; 5xFAD: p = 0.00041, ŠC). During SWRs, more dPCs participated than sPCs (F_(1,87)_ = 12.19, p = 0.00076; 2-way ANOVA, ART; Fig. 3F) in both CT (p = 0.0063, ŠC) and 5xFAD slices (p = 0.00095, ŠC). However, no effect of genotype was observed (F_(1,87)_ = 0.747, p = 0.390), in contrast to the increase observed when considering all PCs together.

We next asked if characteristics of the Ca^2+^ events differed between genotype, as an indirect measure PC firing activity. Averaging across cells from each slice, no differences were found in the frequency (F_(1,87)_ = 0.094, p = 0.759; 2-way ANOVA, log-transformed (LT); Fig. 3G), amplitude (F_(1,87)_ = 0.093, p = 0.761; 2-way ANOVA, LT; Fig. 3H), nor duration of Ca^2+^ transient events (F_(1,87)_ = 0.289, p = 0.593; 2-way ANOVA, LT; Fig. 3I). This suggests that on an individual cell level, PCs exhibit similar activity in 5xFAD mice as compared to controls, with differences only becoming apparent on the ensemble level. Finally, we sought to determine if the cellular composition of PC ensembles during SWR events was altered. Ensemble diversity was assessed by calculating the pairwise Jaccard similarity of cellular participation between all SWR events (Fig. 3J). Additionally, the Jaccard similarity of SWR participation between cells was computed between sPCs/dPCs and all other PCs (Fig. 3J, bottom right). The cumulative distribution functions of all pairwise comparisons were calculated for each slice and averaged across genotype, revealing a reduced degree of similarity in PC ensembles during SWRs in 5xFAD, as compared to control slices (F_(1,4100)_ = 85.6, p < 10^−15^; 2-way ANOVA; Fig. 3K), though with no bins surviving multiple comparisons. This suggests there may be an increased repertoire of ensembles in 5xFAD slices. The similarity between sPCs and all other PCs was reduced (F_(1,3600)_ = 25.2, p = 5.5 × 10^−7^; 2-way ANOVA; Fig. 3L), with similarities between 0.1 – 0.15 surviving multiple comparisons, as was the similarity between dPCs and all other PCs (F_(1,3800)_ = 15.88, p = 6.9 × 10^−5^; 2-way ANOVA; Fig. 3M), suggesting aberrant cell participation may be disrupting ensembles.

### Pyramidal cell spiking is relatively unchanged in 5xFAD mice

To test more directly if pyramidal cell activity was altered, GCaMP6f+ sPCs (n_sPC_ = 13 CT, 9 5xFAD) and dPCs (n_dPC_ = 26 CT, 35 5xFAD) were targeted for loose cell-attached recordings (Fig. 4A). Most cells (n_PC_ = 39 CT, 39 5xFAD) were from 5xFAD;Thy1-GCaMP6f and Thy1-GCaMP6f littermate controls. A small number (n_sPC_ = 1, n_dPC_ = 4) were from 5xFAD;Thy1-GCaMP6f;PV^Cre^/+;tdTom/+ mice, which labeled both excitatory PCs in green and inhibitory PV cells in red. Spikes and bursts (three or more spikes each within 60 ms) were delineated as spontaneous or SWR-coincident (Fig. 4B). Consistent with prior studies (Mizuseki and Buzsáki, 2013), the distribution of spike rates was lognormal for both sPCs and dPCs. Both sPCs (Fig. 4C) and dPCs (Fig. 4D) increased their spike rate during SWRs, with a significant effect of spontaneous/SWR time period (sPCs: F_(1,20)_ = 11.0, p = 0.0034; dPCs: F_(1,59)_ = 18.5, p = 6.4 × 10^−5^; 2-way RM ANOVAs, ART—zero-values preclude LT), with only sPCs/dPCs from 5xFAD mice showing a significant spike rate increase when correcting for multiple comparisons (sPCs: p = 0.272 (CT), p = 0.023 (5xFAD); dPCs: p = 0.085 (CT), p = 0.00013 (5xFAD); Wilcoxon *post hoc*, ŠC). sPCs displayed a significant genotype difference (F_(1,20)_ = 4.64, p = 0.044), and interaction of genotype × period (F_(1,20)_ = 5.59, p = 0.028), though neither the spontaneous nor SWR spike rate survived multiple comparisons (Spont: p = 0.934, SWR: p = 0.301; Mann-Whitney *post hoc*, ŠC). In contrast, dPCs displayed no genotype difference (F_(1,59)_ = 0.914, p = 0.343), nor interaction of genotype × period (F_(1,59)_ = 1.803, p = 0.184).

**Figure 4:**
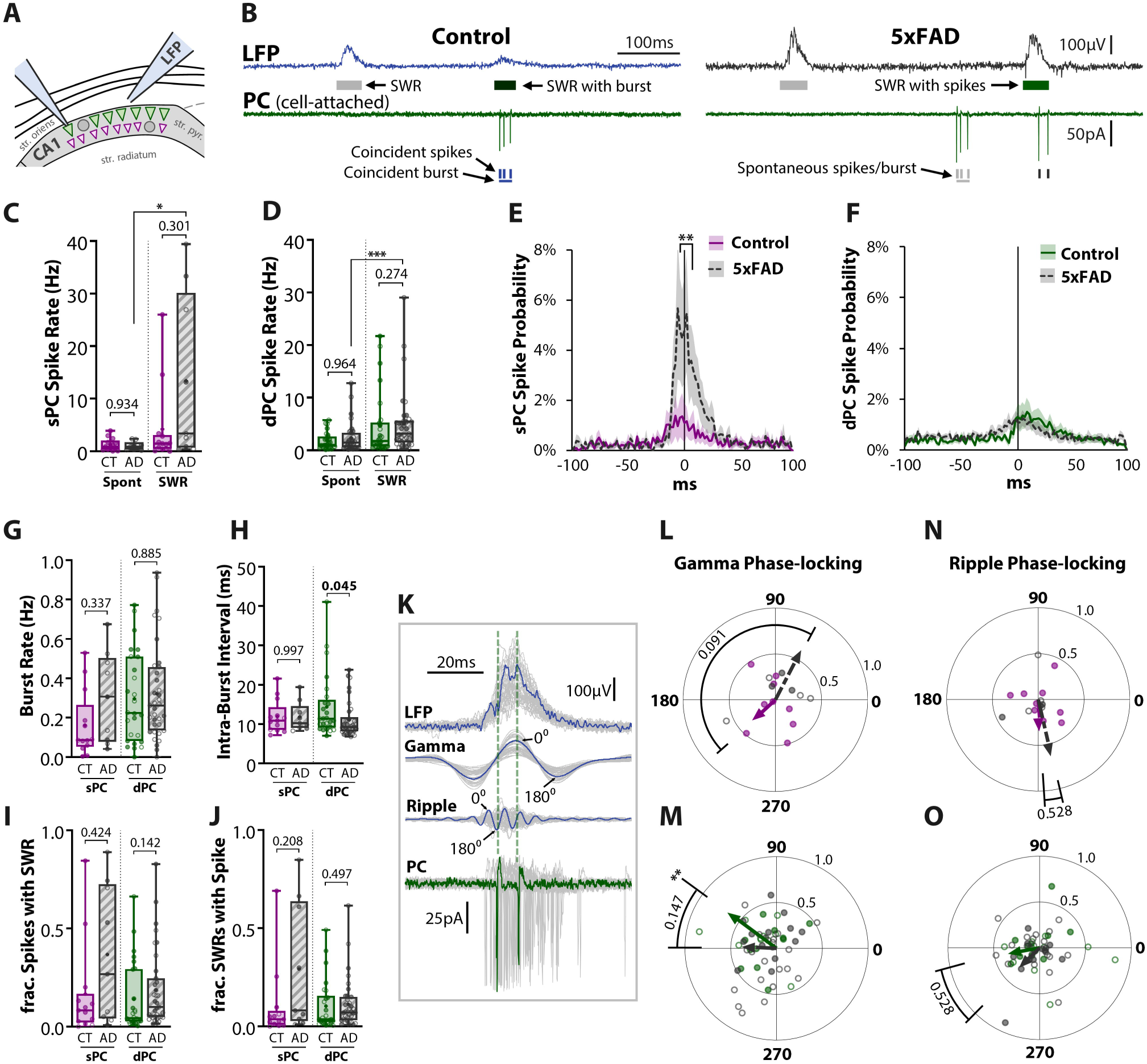
Pyramidal cell spiking is relatively unchanged in 5xFAD mice. ***A***, Diagram of a loose cell-attached recording of spikes from a Thy1-GCaMP6f+ pyramidal cell (PC) paired with LFP recording of SWRs. ***B***, Example traces from 3 mo control (*left*) and 5xFAD (*right*) slice. *Top trace*, LFP with SWR events in raster below. Gray shaded events indicate spontaneous SWR. Green shading indicates SWR coincident with at least one spike. Dark green shading indicates SWR coincident with burst, defined as 3 or more spikes each within 60 ms. *Bottom trace*, Cell-attached recording from PC. Detected spikes and bursts indicated in raster below. Dark shading indicates they are coincident with SWR event, light gray shading indicates spontaneous events. Spike rates were calculated separately during spontaneous and SWR periods for ***C***, superficial PCs (sPC), n_sPC_ = 13 CT, 9 5xFAD, and ***D***, deep PCs (dPCs), n_dPC_ = 26 CT, 35 5xFAD. The peri-SWR spike probability was calculated for ***E***, sPCs and ***F***, dPCs, by binning spikes into 100 2 ms bins and normalizing by total number of SWRs. Dark lines indicate average of all cells within a genotype, shaded regions indicate SEM. 0 ms indicates time of SWR peak. ***G***, The rate of bursts for sPCs and dPCs, defined as at least 3 spikes each within 60 ms. ***H***, Intra-burst interval for sPCs and dPCs, defined as the average time between successive spikes in a burst. ***I***, The fraction of spikes that occurred during SWRs. ***J***, The fraction of SWRs that had one or more spikes. ***K***, Example of SWR (*top trace*), with filtered slow gamma (20-50 Hz) and ripple (120-220 Hz) signals (*middle traces*). The phase of oscillations was set at 0° at peaks and 180° at troughs. *Bottom trace*, cell-attached recording, with spike times marked by vertical dashed lines. ***L-O***, Polar phase plots of phase-locking of spikes to SWR-nested slow gamma (***L-M***) and ripple (***N-O***) for sPCs (***L***,***N***) and dPCs (***M***,***O***). The angles of individual data points represent the average phase of all spikes for a cell. Length from origin (0-1) indicates the degree of phase-locking. A length of 1 signifies perfect phase-locking (every spike at same phase); a length of 0 indicates random or no phase-locking. Lines with arrowheads represent cell average, solid colored = CT, dashed grey = 5xFAD. Asterisks indicate result of Raleigh’s test for non-uniformity, * p < 0.05, ** p < 0.01. p values indicate results of circular mean comparison. For all plots, individual data points represent a cell. Closed circles represent cells from males, open circles females. Box-whisker plots represent non-normal data as Median and IQRs. p-values of pairwise comparisons indicated above brackets.

To examine spike rate in more detail around SWR events, the peri-SWR spike probability was averaged across all cells within each genotype, showing an increased spiking probability in 5xFAD sPCs (F_(1,2000)_ = 37.73, p = 9.8 × 10^−10^; 2-way ANOVA; Fig. 4E), with significant differences when correcting for multiple comparisons from −4 ms to +8 ms around the SWR peak (p < 0.01, ŠC). In contrast, the dPC spike probability did not differ between genotype (F_(1,5900)_ = 2.38, p = 0.123; 2-way ANOVA; Fig. 4F). There was no difference in the rate of spike bursts (3 or more spikes each within 60 ms) between genotype (F_(1,79)_ = 1.00, p = 0.319; 2-way ANOVA, ART; Fig. 4G) or PC type (F_(1,79)_ = 2.37, p = 0.127). There was a tendency for a significant effect of genotype on the intra-burst interval (F_(1,77)_ = 3.26, p = 0.075; 2-way ANOVA, ART; Fig. 4H), with dPCs spiking somewhat faster from 14.4 ± 1.6 to 11.1 ± 0.8 ms (p = 0.045, ŠC) and with no change for sPCs (p = 0.997, ŠC). The percentage of cells with bursts did not differ between genotype (97.6% CT, 96.1% 5xFAD; χ^2^_(3)_= 0.158, p = 0.691; Chi-Squared Test). In particular, 100% of sPCs had at least one burst in both CT and 5xFAD mice, an important consideration given prior work indicating dPCs burst more than sPCs (Mizuseki et al., 2011). As we only targeted cells with visible calcium transients for localization under confocal microscopy, it is possible the population of sPCs and dPCs studied are a more highly active subset of all PCs. We also examined if the fractional participation of PCs in SWRs differed, quantified as the fraction of spikes that occurred during SWRs (Fig. 4I), and the fraction of SWRs that had one or more spikes (Fig. 4J). There was a tendency for an effect of genotype for both endpoints (F_(1,79)_ = 3.90, p = 0.052; F_(1,79)_ = 3.11, p = 0.082; 2-way ANOVA, ART), though with neither cell type surviving multiple comparisons (Fig. 4I-J).

Finally, we examined the phase-locking of spikes in both the slow gamma and ripple ranges (Fig. 4K). Phase-locking of PCs to the trough of ripples has been widely reported *in vivo* (Ylinen et al., 1995; Csicsvari et al., 1999; Le Van Quyen et al., 2008). Additionally, phase-locking of spikes to SWR-nested slow gamma has been reported to be reduced in 5xFAD mice (Iaccarino et al., 2016). We observed that sPCs showed broad phase preference in the gamma range, with averages following the trough at 223° for controls (Z = 0.365, p = 0.706; Raleigh Test), and following the peak at 63° in 5xFAD mice (Z = 1.88, p = 0.154; Raleigh Test), with no significant genotype difference (P = 2.86, p = 0.091, Circular-Median Test; Fig. 4L). In contrast, control dPCs were significantly phase-locked at 143° (110°-177° 95%; Z = 5.79, p = 0.0023; Raleigh Test), while 5xFAD dPCs showed a broader phase preference with a mean of 177° (110°-245° 95%; Z = 2.25, p = 0.105; Raleigh Test), and no significant genotype difference (P = 0.147, p = 0.701, Circular-Median Test; Fig. 4M). In the ripple range cells displayed broad phase preference, with neither sPCs Z = 0.355, p = 0.712; Z = 1.58, p = 0.212; Raleigh Test; Fig. 4N) nor dPCs (Z = 0.981, p = 0.340; 5xFAD: Z = 1.09, p = 0.339; Raleigh Test; Fig. 4O) significantly phase-locked, nor different between genotypes (sPC: P = 0.254, p = 0.614; dPC: P = 0.528, p = 0.467; Circular-Median Test). However, in control dPCs, the mean phase at the trough of 191° more closely matches prior *in vivo* findings than for 5xFAD mice, with a mean phase of 224° following the trough (Fig. 4O). These results indicate that spike-phase coupling may be impaired in 5xFAD mice, though the lack of significant ripple phase-locking for either genotype suggests there are limitations to this analysis in our slice preparation, as this finding differs from the robust ripple phase-locking observed *in vivo* (Ylinen et al., 1995; Csicsvari et al., 1999; Le Van Quyen et al., 2008). One difference between our study and prior *in vivo* recordings is that the spikes and LFP were recorded from different electrodes, while *in vivo* LFP and spikes are typically recorded from the same set of channels. As ripples are highly localized events, the distance between electrodes may confound spatial phase locking. Taken together with the Ca^2+^ imaging data, these results indicate relatively minor alterations to the spiking activity of PCs in 5xFAD slices. However, the large variability in spiking rate may mask small differences in activity.

### Pyramidal cells receive increased synaptic E/I ratio

Following the cell-attached recording of PC spiking activity, the electrode was replaced with one containing a Cesium-based internal solution and the same cell targeted for a whole-cell voltage-clamp recording to detect excitatory and inhibitory post-synaptic currents (EPSCs and IPSCs) at −70 mV (n_sPC_ = 7 CT, 5 5xFAD; n_dPC_ = 12 CT, 9 5xFAD; Fig. 5A-B) and 0 mV (n_sPC_ = 7 CT, 5 5xFAD; n_dPC_ = 14 CT, 11 5xFAD; Fig. 5C-D), respectively. Events were sorted as spontaneous (sEPSCs/sIPSCs) or SWR-coincident (swrEPSCs/swrIPSCs). sEPSC frequency and amplitude were unchanged across genotype (Freq: F_(1,29)_ = 0.296, p = 0.591; Fig. 5E; Amp: F_(1,29)_ = 2.26, p = 0.143; Fig. 5F; 2-way ANOVA, LT), as were the kinetics of sEPSCs (Rise tau: F_(1,29)_ = 0.540, p = 0.468; Fig. 5G; Decay tau: F_(1,29)_ = 0.268, p = 0.608; Fig. 5H; 2-way ANOVA). The excitatory charge during spontaneous periods (sEPSQ) was also unchanged across genotype (F_(1,29)_ = 0.835, p = 0.369; 2-way ANOVA, LT; Fig. 5I). However, there was a significant effect of genotype on the excitatory charge during SWRs (swrEPSQ) (F_(1,29)_ = 6.56, p = 0.016; 2-way ANOVA, LT; Fig. 5J), with dPCs seeing a 114 ± 41% increase (p = 0.043, ŠC), while sPCs were not significantly different (p = 0.368, ŠC). Considering we observed that SWRs in 5xFAD mice were larger in amplitude with more total PCs active, this was not an altogether surprising result. This increase indicates that the enlarged SWRs were accompanied with increased excitatory synaptic activity, likely originating from the CA3 region. The lack of any increase in spontaneous excitatory activity is also consistent with our cell-attached results showing that at least locally in CA1, both sPCs and dPCs had no changes in firing rate during spontaneous periods (Fig. 4C-D). Together these results indicate that spontaneous excitatory synaptic input is unchanged, but increased during SWRs in 5xFAD mice, particularly for dPCs, consistent with expectations from our LFP experiments.

**Figure 5:**
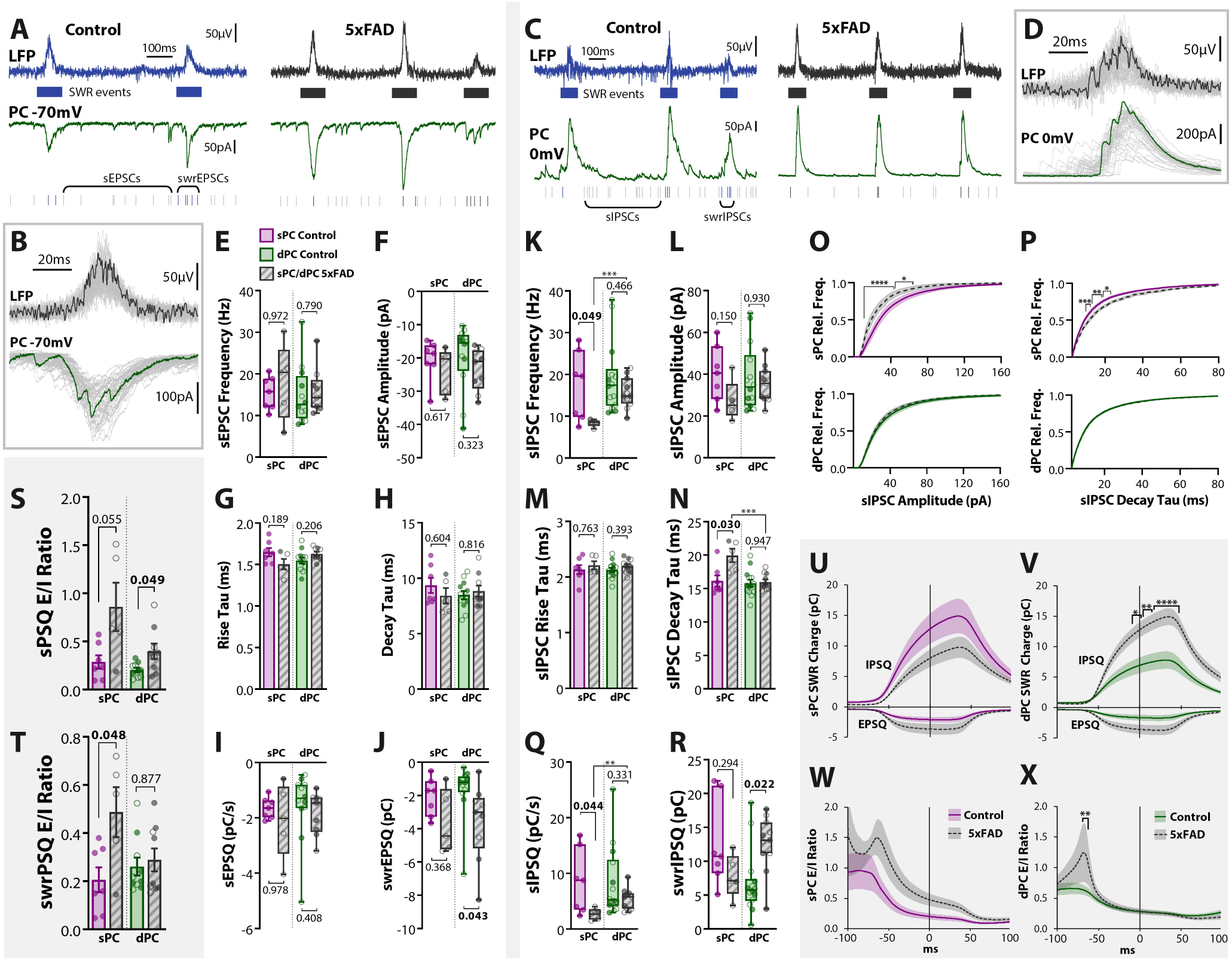
Pyramidal cells receive increased synaptic E/I ratio. ***A***, Example traces of LFP + whole-cell recording voltage-clamped at −70 mV in Thy1-GCaMP6f littermate controls and 5xFAD/+;Thy1-GCaMP6f mice. Excitatory post-synaptic potentials (EPSCs) were sorted as spontaneous (sEPSCs) or SWR-coincident (swrEPSCs). ***B***, Example swrEPSCs. ***C***, Example traces of LFP + whole-cell recording voltage-clamped at 0 mV to detect sIPSCs and swrIPSCs. ***D***, Example swrIPSCs. ***E-H***, Summary plots of sEPSC frequency (***E***), amplitude (***F***), rise tau (***G***), and decay tau (***H***) for sPCs and dPCs in control and 5xFAD mice (n_sPC_ = 7 CT, 5 5xFAD; n_dPC_ = 12 CT, 9 5xFAD). ***I***, sEPSQ = normalized spontaneous excitatory charge (integrated current per unit time). ***J***, swrEPSQ = total excitatory charge during SWRs, integrated over a 100 ms window centered around the SWR peak. ***K-N***, Summary plots of sIPSC frequency (***K***), amplitude (***L***), rise tau (***M***), and decay tau (***N***) (n_sPC_ = 7 CT, 5 5xFAD; n_dPC_ = 14 CT, 11 5xFAD). ***O-P***, Cumulative distribution functions, averaged over all cells for sIPSC amplitude (***O***) and decay tau (***P***), both for sPCs (*top*) and dPCs (*bottom*). Dark lines represent cell average, shaded region represents SEM. ***Q***, sIPSQ = normalized spontaneous inhibitory charge (integrated current per unit time). ***R***, swrIPSQ = total inhibitory charge during SWRs. ***S***, Spontaneous synaptic E/I ratio of normalized charge for each cell (sEPSQ/sIPSQ). ***T***, Synaptic E/I ratio for 100 ms window centered around SWR peak (swrEPSQ/swrIPSQ). ***U-V***, During SWRs, the charge for each cell (EPSQ and IPSQ) was calculated in 2 ms bins by integrating the current in a sliding 100 ms window centered around that bin, for both sPCs (***U***) and dPCs (***V***). Summary data in ***J*** and ***R*** thus represent these curves at the y-intercept. Asterisks indicate regions that survived Šidák’s multiple comparisons * < 0.05, ** < 0.01, *** < 0.01. ***W-X***, the ratios of curves in ***U-V*** yield the synaptic E/I ratio during SWRs. Summary data in ***T*** represents these curves at the y-intercept. For all plots, individual data points represent a cell. Closed circles represent cells from males, open circles females. Box-whisker plots represent non-normal data with Median and IQRs. Bar plots represent normal data with Mean ± SEM. p-values of pairwise comparisons indicated above brackets.

In contrast, for sIPSCs, there was an effect of genotype on the frequency (F_(1,33)_ = 9.38, p = 0.0043; 2-way ANOVA, LT; Fig. 5K), as well as an effect of PC type (F_(1,33)_ = 8.19, p = 0.0073), with sPCs seeing a preferential 52 ± 10% decrease (p = 0.049, ŠC), while dPCs were unchanged (p = 0.466, ŠC). There was a tendency for an effect of genotype on the amplitude of sIPSCs (F_(1,33)_ = 3.40, p = 0.074; 2-way ANOVA, LT; Fig. 5L), and no effect for the rise tau (F_(1,33)_ = 1.73, p = 0.197; 2-way ANOVA; Fig. 5M). However the decay tau of sIPSCs saw an effect of genotype (F_(1,33)_ = 9.36, p = 0.0044; 2-way ANOVA; Fig. 5N), cell type (F_(1,33)_ = 10.7, p = 0.0025), and interaction of genotype × cell type (F_(1,33)_ = 7.61, p = 0.0094), with sPCs from 5xFAD mice displaying a 5 ms longer decay than control sPCs (p = 0.030, ŠC) and dPCs (p = 0.00072, ŠC). This specific alteration prompted us to examine in more detail the distribution of sIPSCs. For each cell, the cumulative distribution function of all events was calculated and averaged across cells. This analysis revealed for sPCs a significantly lower sIPSC amplitude (F_(1,1000)_ = 39.8, p = 4.2 × 10^−15^; 2-way ANOVA; Fig. 5O, upper) and greater decay tau (F_(1,1000)_ = 469, p < 10^−15^; 2-way ANOVA; Fig. 5P, upper), with events of amplitude 10 – 65 pA and decay tau 9 – 22 ms surviving multiple comparisons. This suggests that for sPCs, fast and high amplitude inhibitory input, typically attributed to fast-spiking cells, is preferentially reduced. In contrast, dPCs saw no significant range of bins survive multiple comparisons (Fig. O-P, lower). This analysis was performed for all sEPSC and sIPSC end-points, and no other end-points saw significant genotype differences surviving multiple comparisons.

The normalized inhibitory charge during spontaneous periods (sIPSQ) was also preferentially reduced for 5xFAD sPCs (Genotype Effect: F_(1,33)_ = 10.3, p = 0.0029; 2-way ANOVA, LT; sPC: p = 0.044, dPC: p = 0.331, ŠC; Fig. 5Q). During SWRs, the total inhibitory charge (swrIPSQ) was differentially altered between cell types, with a significant interaction of genotype and cell type (F_(1,33)_ = 7.14, p = 0.0116; 2-way ANOVA, LT; Fig. 5R), a non-significant reduction for sPCs (p = 0.294, ŠC) and a significant increase for dPCs (p = 0.022, ŠC). These results suggest a selective impairment in inhibition in sPCs that is not seen in dPCs. Despite the observed increase in SWR activity in 5xFAD mice, the inhibition did not scale proportionally with the increased excitation, thus shifting the synaptic E/I balance. We assessed this for both spontaneous and SWR-driven currents. During spontaneous periods, there was a significant effect of genotype on the E/I ratio, defined as the ratio of sEPSQ/sIPSQ (F_(1,27)_ = 14.7, p = 0.00069; 2-way ANOVA; Fig. S), as well as an effect of cell type (F_(1,27)_ = 7.39, p = 0.0113). Both sPCs and dPCs saw an increase (sPC: p = 0.055, dPC: p = 0.049, ŠC). During SWRs, the synaptic E/I ratio (swrEPSQ/swrIPSQ) was affected by both genotype (F_(1,27)_ = 7.59, p = 0.0104; 2-way ANOVA; Fig. R) and a genotype x cell type interaction (F_(1,27)_ = 5.12, p = 0.0319), but only in sPCs was there an increase (sPC: p = 0.048, dPC: p = 0.877). To examine the temporal progression of synaptic input during SWRs, we calculated the EPSQ and IPSQ in a sliding 100 ms window across the SWR peak (Fig. 5U-V), from which a time course of the synaptic E/I ratio could be determined (Fig. 5W-X). While for both the EPSQ and IPSQ there was a significant effect of genotype for sPCs (EPSQ: F_(1,1000)_ = 184, p < 10^−15^; IPSQ: F_(1,1000)_ = 129, p < 10^−15^; 2-way ANOVA; Fig. 5U) and dPCs (EPSQ: F_(1,1900)_ = 295, p < 10^−15^; IPSQ: F_(1,2300)_ = 708, p < 10^−15^; 2-way ANOVA; Fig. 5V), only for the IPSQ in dPCs did individual bins from −18 to +62 ms (relative to the SWR peak) survive multiple comparisons. The ratio of the EPSQ/IPSQ time course revealed a significant effect of genotype for both sPCs (F_(1,1000)_ = 159, p < 10^−15^; 2-way ANOVA; Fig. 5W) and dPCs (F_(1,1800)_ = 24.3, p = 9.1 × 10^−7^; 2-way ANOVA; Fig. 5X). For both sPCs and dPCs there was an early peak in the synaptic E/I ratio for 5xFAD mice that was not present in controls (Fig. 5W-X), which can be attributed to a build-up in excitation with a delayed increase in inhibition. Only for dPCs did individual bins from −70 to −60 ms survive multiple comparisons, in part likely due to the greater sampling of dPCs.

### PV basket cells have selectively reduced spiking

While there are numerous inhibitory cell sub-types in the CA1 region that could underlie a shift in E/I synaptic input to PCs (Pelkey et al., 2017), we focused on parvalbumin-expressing (PV) interneurons, as they are the most highly active during SWR events (Somogyi et al., 2014). We performed cell-attached recordings in 5xFAD/+;PV^Cre^/+;tdTom/+ and PV^Cre^/+;tdTom/+ littermate controls (n_cell_ = 13 CT, 18 5xFAD) (Fig. 6A). Cells were also recorded from 5xFAD;Thy1-GCaMP6f;PV^Cre^/+;tdTom/+ and Thy1-GCaMP6f;PV^Cre^/+;tdTom/+ littermate controls (n_cell_ = 11 CT, 14 5xFAD), which were pooled together. One complication with the PV cell population is that there are at least three distinct sub-types within CA1 *str. pyr.*, which vary in function and axonal target: basket cells (PVBCs), which target perisomatic regions of PCs, bistratified cells (PVBSCs), which target both apical and basal dendrites of PCs, and axo-axonic cells (PVAACs), which selectively target the axon initial segment (AIS) (Fig. 6B). To distinguish these, we morphologically reconstructed the cells *post hoc* after filling with biocytin in whole-cell configuration (Fig. 6C) and sorted them by axonal target. Of the total reconstructed cells (n_cell_ = 24 CT, 32 5xFAD), PVBSCs (Fig. 6C.2) were easily distinguished from both PVBCs (Fig. 6C.1) and PVAACs (Fig. 6C.3), with their axonal arbor avoiding *str. pyramidale* (n_PVBSC_ = 5 CT, 10 5xFAD). While PVBCs and PVAACs have overlapping axonal targets, some PVBCs were easily distinguished with axonal terminals directly targeting PC somas as visualized through the PV^Cre^-tdTom and/or Thy1-GCaMP6f fluorescence (n_PVBC_ = 9 CT, 9 5xFAD). Likewise, some PVAACs exhibited the characteristic “chandelier” phenotype and lack of somatic targeting (n_PVAAC_ = 4 CT, 4 5xFAD). However, there were some cells with ambiguous PVBC/PVAAC morphology based solely on axonal targets (n_cell_ = 6 CT, 9 5xFAD). To sort these cells, we examined the spike rate. Both PVBCs and PVBSCs are known to strongly increase their spike rate during SWRs (Lapray et al., 2012; Katona et al., 2014), while the PVAAC spike rate does not increase (Viney et al., 2013). Thus, for cells with ambiguous PVBC/PVAAC morphology, those with a spike rate increase during SWRs (53 ± 11 Hz) were sorted as putative PVBCs, while those with no increase (0.6 ± 0.6 Hz) were sorted as putative PVAACs. In a subset of 14 slices (5 CT, 9 5xFAD from n_mice_ = 4 CT, 5 5xFAD), we additionally stained for Ankyrin G, which labels the AIS, and confirmed colocalization with two putative PVAACs (Fig. 6C.4). Based on this sorting methodology, we identified a total of n_PVBC_ = 13 CT, 16 5xFAD, n_PVBSC_ = 5 CT, 10 5xFAD, and n_PVAAC_ = 6 CT, 6 5xFAD. The proportion of cells did not significantly differ from prior published findings of 60% PVBC, 25% PVBSC, 15% PVAAC in CA1 *str. pyr.* (χ^2^_(2)_ = 0.754, p = 0.686) (Baude et al., 2007), and did not significantly differ between genotype (χ^2^ _(2)_ = 1.09, p = 0.580; Fig. 6D).

**Figure 6:**
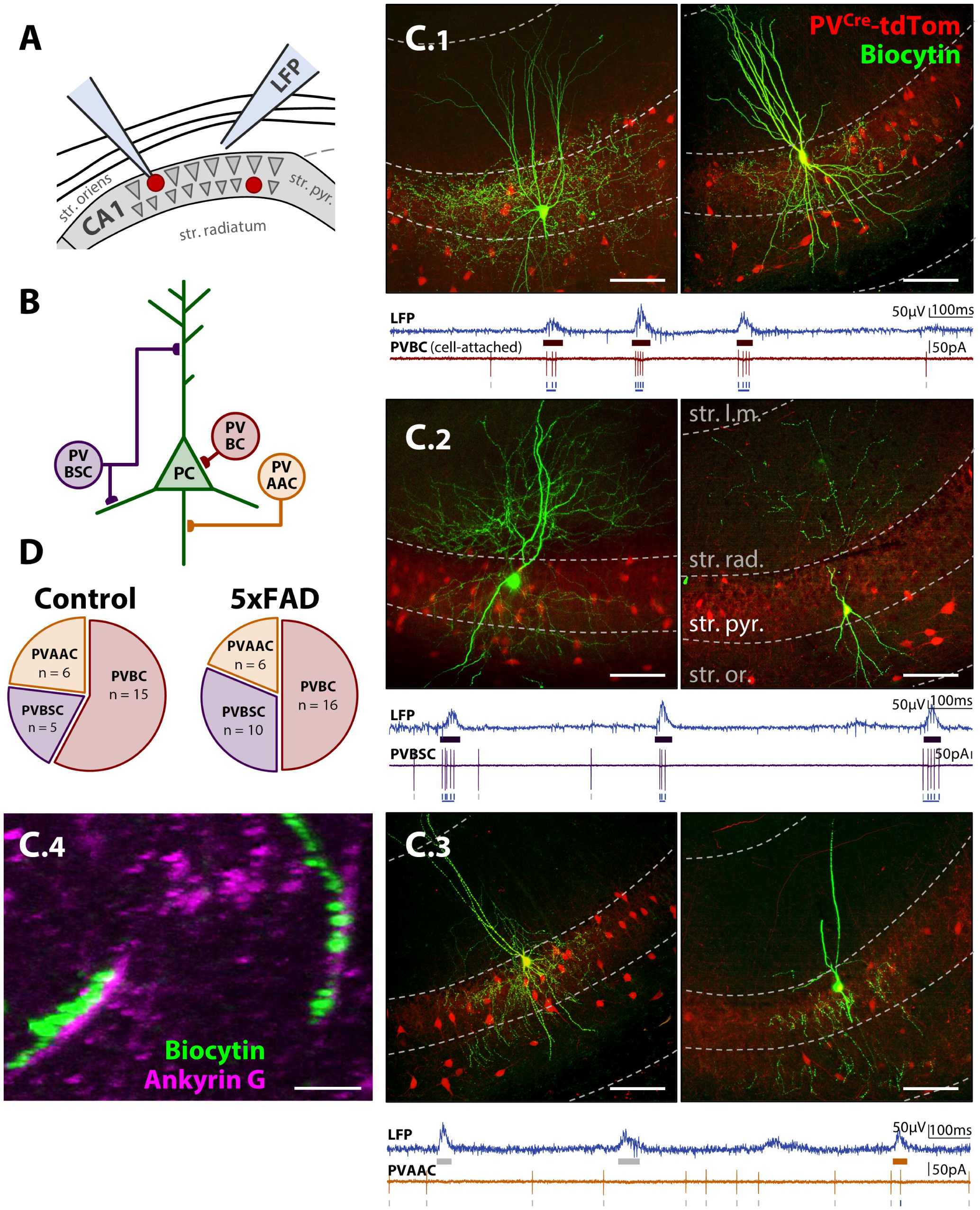
PV cells in CA1 delineate into basket (PVBCs), bistratified (PVBSCs), and axo-axonic (PVAACs) cells. ***A***, Diagram of LFP + PV cell recording in 3 mo 5xFAD/+; PV^Cre^/+;tdTom/+ and PV^Cre^/+;tdTom/+ littermate controls. ***B***, Diagram of PV cell subtypes and their axonal targets in CA1 *str. pyr.* Examples of ***C.1***, PVBCs, ***C.2***, PVBSCs, and ***C.3***, PVAACs. Green fluorescence indicates avidin-fluorescein bound to the biocyin in the filled PV cell. Red fluorescence indicates PV^Cre^/+;tdTom/+ expression. Dashed grey lines approximately distinguish layers of hippocampus, *str. l.m.* = *stratum lacunosum-moleculare, str. rad.* = *stratum radiatum, str. pyr.* = *stratum pyramidale, str. or.* = *stratum oriens*. Notice that *str. pyr.* is dimly red due to the expression of other non-filled PVBCs targeting perisomatic regions of PCs. Scale bars = 100 μm. *Bottom traces*, Examples of LFP and cell-attached recording of spikes, which was used to aid in distinguishing PVBCs and PVAACs. PVAACs are unique in that they reduce their firing during SWRs. ***C.4***, Ankyrin G immunostaining showing co-localization of synaptic boutons with AIS of PCs for identified PVAAC. Scale bar = 10 μm. ***D***, Proportions of sorted cells in each genotype.

Based on these delineations, we performed PV cell-attached + LFP recordings in control and 5xFAD mice for the identified populations of PVBCs (Fig. 7A.1), PVBSCs (Fig. 7A.2) and PVAACs (Fig. 7A.3). We observed an effect of genotype on PVBC spike rate (F_(1,27)_ = 10.4, p = 0.0033; 2-way RM ANOVA, ART; Fig. 7B.1), with a selective and robust reduction during SWR periods, from 62.9 ± 10.6 Hz to 34.0 ± 6.2 Hz (p = 0.044; Mann-Whitney *post hoc*, ŠC), whereas the spontaneous spike rate did not significantly differ (p = 0.209; Mann-Whitney *post hoc*, ŠC). In contrast, there was no effect of genotype on the spike rate of PVBSCs (F_(1,13)_ = 1.35, p = 0.267; 2-way RM ANOVA, ART; Fig. 7B.2) or PVAACs (F_(1,10)_ = 3.83, p = 0.079; 2-way RM ANOVA, ART; Fig. 7B.3). As expected from the cell-sorting methodology, spike rates increased during SWRS for PVBCs, with an effect of period (F_(1,27)_ = 179, p = 1.9 × 10^−13^; Fig. 7B.1), and increases of 5.6 ± 1.2 fold in control (p = 0.00049; Wilcoxon *post hoc*, ŠC) and 5.9 ± 2.4 fold in 5xFAD mice (p = 0.00012; Wilcoxon *post hoc*, ŠC). Additionally, there was an interaction of genotype × period (F_(1,27)_ = 16.8, p = 0.00034), indicating that PVBC modulation of spiking during SWRs differed between genotype. Similarly, there was an effect of period for PVBSCs (F_(1,13)_ = 30.6, p = 9.7 × 10^−5^; Fig. 7B.2), with a non-significant 4.4 ± 1.2 fold increase in control (p = 0.121; Wilcoxon *post hoc*, ŠC) and a 6.8 ± 5.0 fold increase in 5xFAD mice (p = 0.019; Wilcoxon *post hoc*, ŠC), and no interaction of genotype × period (F_(1,13)_ = 2.56, p = 0.133). While PVAACs did see an effect of period (F_(1,10)_ = 16.7, p = 0.0022; Fig. 7B.3), neither in control nor 5xFAD mice was the increase significant (CT: 1.0 ± 0.4 fold, p = 0.527, 5xFAD: 2.4 ± 0.8 fold, p = 0.062; Wilcoxon *post hoc*, ŠC), nor was there an interaction of genotype × period (F_(1,10)_ = 3.21, p = 0.104). However, the low number of PVAACs recorded from may mask small alterations in this cell population.

**Figure 7:**
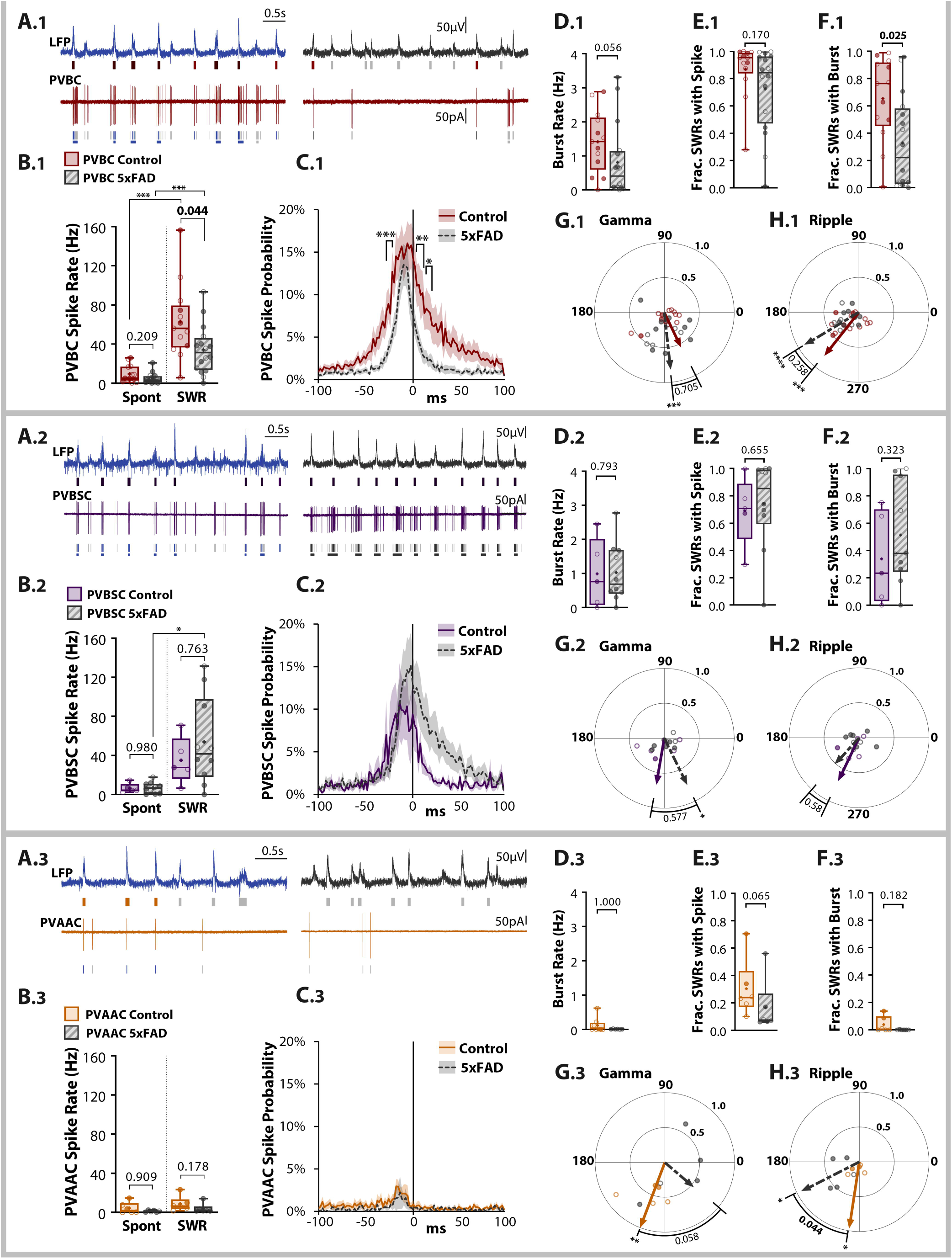
PV basket cells have selectively reduced spiking. Spiking data for PVBCs (panel sub-heading ***1***), PVBSCs (panel sub-heading ***2***), and PVAACs (panel sub-heading ***3***). ***A***, Example LFP and cell-attached traces for CT (*left*) and 5xFAD (*right*), for a PVBC (***A.1***), PVBSC (***A.2***), and PVAAC (***A.3***). Spike and SWR events are color-coded as in Fig. 4B. ***B***, Summary spike rate data for PVBCs (n_PVBC_ = 13 CT, 16 5xFAD, ***B.1***), PVBSCs (n_PVBSC_ = 5 CT, 10 5xFAD, ***B.2***), and PVAACs (n_PVAAC_ = 6 CT, 6 5xFAD, ***B.3***) during spontaneous and SWR periods. ***C***, Peri-SWR spike probability. Asterisks indicate regions surviving Šidák’s multiple comparisons correction. ***D***, Burst rate, defined as 3 or more spikes each within 40 ms. ***E***, The fraction of SWRs with one or more spike. ***F***, The fraction of SWRs with a burst. ***G-H***, Polar phase plots of phase-locking of spikes to SWR-nested slow gamma (***G***) and ripple (***H***). The angles of individual data points represent the average phase of all spikes for a cell. Length from origin (0-1) indicates the degree of phase-locking. Lines with arrowheads represent cell average, solid colored = CT, dashed grey = 5xFAD. Asterisks indicate result of Raleigh’s test for non-uniformity, p values indicate results of Watson-Williams or circular mean test. * < 0.05, ** < 0.01, *** < 0.001. For all plots, individual data points represent a cell. Closed circles represent cells from males, open circles females. Box-whisker plots represent non-normal data with Median and IQRs. p-values indicated above brackets.

The peri-SWR spike probability revealed that in 5xFAD mice, PVBCs spikes fell in a significantly narrower window (Fig. 7C.1), with a reduced full-width half-maximum value from 44 ± 19 ms to 18 ± 5 ms, and a significant effect of genotype when analyzed via 2-way ANOVA (F_(1,2700)_ = 267, p < 10^−15^), with bins from −20 to −16 ms and 0 to +20 ms relative to the SWR peak surviving multiple comparisons. Intriguingly, this narrower window of spiking in 5xFAD mice was accompanied by shorter duration SWRs (Fig. 2D), suggesting the activity of these cells is critical for normal ripple progression. In contrast, PVBSCs appeared to increase their firing after the SWR peak in 5xFAD mice (Fig. 7C.2), perhaps playing a compensatory role, with a significant effect of genotype (F_(1,1300)_ = 33.9, p = 7.3 × 10^−9^; 2-way ANOVA). PVAACs appeared to have decreased probability of spiking in 5xFAD mice, particularly before the SWR peak (Fig. 7C.3), with a significant effect of genotype (F_(1,1000)_ = 30.15, p = 5.1 × 10^−8^; 2-way ANOVA). However, unlike PVBCs, neither PVBSCs nor PVAACs showed significant genotype differences when corrected for multiple comparisons (Fig. 7C).

Considering the rate of bursts, defined as three or more spikes each within 40 ms, there was a tendency for a reduction for PVBCs (U = 60, p = 0.056; Mann-Whitney; Fig. 7D.1) and no change for PVBSCs (U = 22.5, p = 0.793; Mann-Whitney; Fig. 7D.2) or PVAACs (U = 24, p > 0.999; Mann-Whitney; Fig. 7D.3). No differences were observed in the proportion of cells that exhibited bursts for PVBCs (100% CT, 93.8% 5xFAD; χ^2^_(1)_ = 0.842, p = 0.359; χ^2^ Test), PVBSCs (80.0% CT, 90.0% 5xFAD; χ^2^_(1)_ = 0.288, p = 0.591), or PVAACs (50.0% CT, 83.3% 5xFAD; χ^2^_(1)_ = 1.500, p = 0.221). The fraction of SWRs that coincided with a PV spike was no different between genotypes for PVBCs (U = 72, p = 0.170; Mann-Whitney; Fig. 7E.1), PVBSCs (U = 21, p = 0.655; Mann-Whitney; Fig. 7E.2), or PVAACs (U = 6, p = 0.065; Mann-Whitney; Fig. 7E.3). However in PVBCs, the fraction of SWRs that coincided with a PV burst was significantly reduced from 65.4 ± 8.5% to 32.9 ± 8.5% (U = 53, p = 0.025; Mann-Whitney; Fig. 7F.1), while there was no genotype difference for PVBSCs (U = 16.5, p = 0.323; Mann-Whitney; Fig. 7F.2) or PVAACs (U = 10.5, p = 0.182; Mann-Whitney; Fig. 7F.3).

We also examined the spike phase-locking of these three PV cell types, as these have been carefully studied *in vivo* for theta (8-12 Hz, during mobility) and ripple oscillations (Varga et al., 2014). The precise temporal ordering of PV cell sub-types during network oscillations has been proposed to play a critical role in the spatiotemporal control of PCs. During ripples, PVBCs have been observed to fire just after the trough of the ripple, followed by PVBSCs and then PVAACs. Less studied is the phase-locking of PV cells during SWR-nested slow gamma oscillations, which we examined as we observed alterations in this endpoint for the PC population (Fig. 4M). We found significant gamma phase-locking of PVBCs for 5xFAD mice at 273° (247°-298° 95%; Z = 7.94, p = 0.00012; Raleigh’s test), with broader phase preference in control mice (Z = 2.48, p = 0.082; Raleigh’s test), and with no difference in median phase angle (P = 0.144, p = 0.705; Circular median test; Fig. 7G.1). PVBSCs were similarly gamma phase-locked for 5xFAD mice at 292° (249°-336° 95%; Z = 3.54, p = 0.024; Raleigh’s test), with broader phase preference in control mice (Z = 1.57, p = 0.216; Raleigh’s test), and with no difference in median phase angle (P = 0.311, p = 0.577; Circular median test; Fig. 7G.2). In contrast, in PVAACs we observed phase-locking in control mice at 246° (224°-368° 95%; Z = 4.43, p = 0.005; Raleigh’s test), with broader phase preference in 5xFAD mice (Z = 0.913, p = 0.421; Raleigh’s test), and with a tendency for a difference in median phase angle (P = 3.60, p = 0.058; Circular median test; Fig. 7G.3). While the precise significance of gamma phase-locking has yet to be demonstrated, these genotype differences point to a temporal disordering of PV cell inhibition.

Within the ripple range, in PVBCs, there was a significant phase-locking in control mice at 228° (198°-259° 95%; Z = 6.01, p = 0.0014; Raleigh’s test), and in 5xFAD mice at 208° (186°-231° 95%; Z = 9.35, p = 1.7 × 10^−5^; Raleigh’s test), with no significant genotype difference (F_(1,27)_ = 1.34, p = 0.258; Watson-Williams; Fig. 7H.1). These values are in line with prior *in vivo* studies, where 0° in our study was defined as the peak of the ripple cycle and 180° as the trough. PVBSCs exhibited more varied phase-locking in the ripple range, although the average angles are in line with *in vivo* studies (CT: 241°, Z = 1.63, p = 0.202; 5xFAD: 225°, Z = 1.33, p = 0.270; Raleigh’s test; Fig. 7H.2). PVAACs exhibited ripple phase preference for control mice at 257° (219°-296° 95%; Z = 3.75, p = 0.0015; Raleigh’s test), in line with *in vivo* studies. However, in 5xFAD mice, PVAACs spiked earlier, at 204° (164°-245° 95%; Z = 3.54, p = 0.0020; Raleigh’s test), with a significant genotype difference in mean phase angle (F_(1,9)_ = 5.68, p = 0.044; Watson-Williams; Fig. 7H.3). Although we observed no significant change in PVAAC spike rate, this disruption in temporal ordering may still have downstream network consequences.

### PV basket cells have selective decrease in excitatory synaptic drive and decreased E/I ratio

Following cell-attached recording of spiking activity, the electrode was replaced with a Cesium internal and the same PV cell was targeted for a whole-cell voltage-clamp recording. EPSCs were recorded at −70 mV for PVBCs (n_PVBC_ = 12 CT, 16 5xFAD; Fig. 8A.1), PVBSCs (n_PVBSC_ = 4 CT, 6 5xFAD; Fig. 8A.2), and PVAACs (n_PVAAC_ = 5 CT, 6 5xFAD; Fig. 8A.3). None of the three PV cell subtypes had any change in spontaneous sEPSC frequency (PVBC: t_(26)_ = 1.30, p = 0.206; PVBSC: t_(8)_ = 0.990, p = 0.351; PVAAC: t_(9)_ = 0.129, p = 0.900; unpaired t-tests), or amplitude (PVBC: t_(26)_ = 0.545, p = 0.591; PVBSC: t_(8)_ = 0.058, p = 0.956; PVAAC: t_(9)_ = 2.11, p = 0.064; unpaired t-tests; Fig. 8B.1-3). The kinetics of sEPSCs in 5xFAD PVBCs were altered however, with a similar rise tau (t_(26)_ = 0.307, p = 0.761; unpaired t-test), but a reduction in the decay tau (t_(26)_ = 2.41, p = 0.024; unpaired t-test; Fig. 8C.1). There were no changes to sEPSC kinetics in PVBSCs (Rise: t_(8)_ = 0.671, p = 0.521; Decay: t_(8)_ = 1.02, p = 0.340; Fig. 8C.2) or PVAACs (Rise: t_(9)_ = 0.857, p = 0.414; Decay: t_(9)_ = 1.79, p = 0.107; Fig. 8C.3). To examine the decreased sEPSC decay tau in PVBCs, the cumulative distribution function of all sEPSCs was calculated and averaged across cells, revealing an effect of genotype (F_(1,2500)_ = 2165, p < 10^−15^; 2-way ANOVA; Fig. 8D), with events of decay tau 1 – 7 ms surviving multiple comparisons. This decrease in decay tau was somewhat unexpected, considering that in PV cells, neuronal pentraxins are associated with an acquisition of GluA4 subunits which speeds up the kinetics of AMPA receptor mediated EPSCs (Pelkey et al., 2015). Both pentraxins and GluA4 are selectively reduced in human Alzheimer’s patients (Xiao et al., 2017), suggesting that in AD, a longer decay of AMPA-mediated EPSCs may underlie PV cell dysfunction. Our results indicate that in 3 mo 5xFAD mice, this does not appear to be a prominent mechanism.

**Figure 8:**
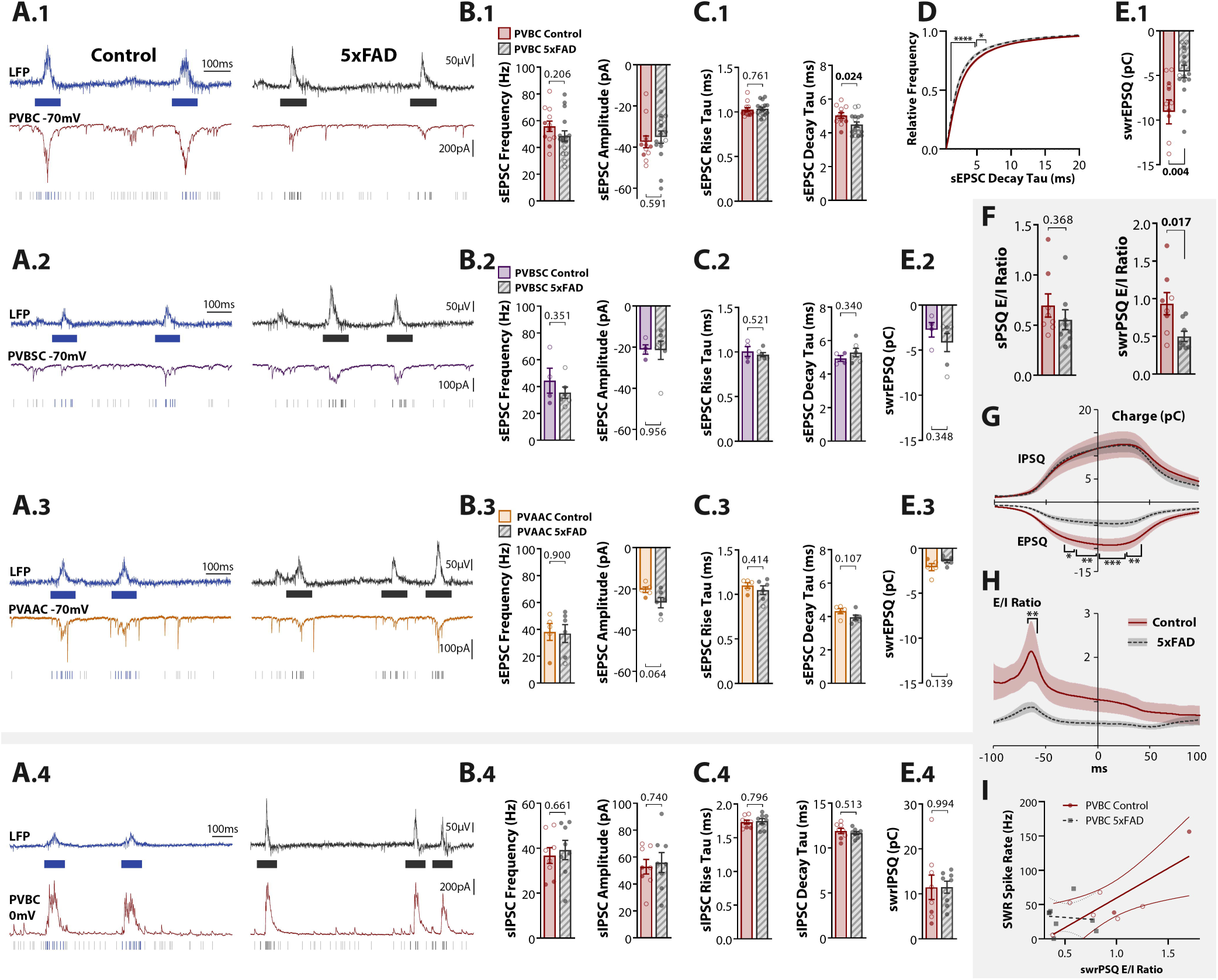
PV basket cells have selective decrease in excitatory synaptic input and decreased E/I ratio. Whole-cell data for PVBC EPSCs (panel sub-heading ***1***), PVBSC EPSCs (panel sub-heading ***2***), PVAAC EPSCs (panel sub-heading ***3***), and PVBC IPSCs (panel sub-heading ***4***). ***A***, Example recordings of LFP + whole-cell PV cell recording in 3 mo 5xFAD/+; PV^Cre^/+;tdTom/+ and PV^Cre^/+;tdTom/+ littermate controls, recording synaptic input during spontaneous and SWR period. EPSCs were recorded at −70 mV in PVBCs (***A.1***), PVBSCs (***A.2***), and PVAACs (***A.3***), and IPSCs recorded at 0 mV in a subset of PVBCs (***A.4***). ***B***, Summary data for spontaneous PSC frequency and amplitude for PVBC sEPSCs (n_PVBC_ = 12 CT, 16 5xFAD, ***B.1***), PVBSC sEPSCs (n_PVBSC_ = 4 CT, 6 5xFAD, ***B.2***), PVAAC sEPSCs (n_PVAAC_ = 5 CT, 6 5xFAD, ***B.3***), and PVBC sIPSCs (n_PVBC_ = 8 CT, 8 5xFAD, ***B.4***). ***C***, Spontaneous PSC kinetics: rise and decay tau. ***D***, Cumulative distribution function of PVBC decay tau. Dark lines represent cell average, shaded region represents SEM. ***E***, swrEPSQ and swrIPSQ = total excitatory and inhibitory charge during SWRs, integrated over a 100 ms window centered around the SWR peak. ***F***, *left*, PVBC spontaneous synaptic E/I ratio of normalized charge (sEPSQ/sIPSQ). ***F***, *right*, PVBC synaptic E/I ratio for 100 ms window centered around SWR peak (swrEPSQ/swrIPSQ). ***G***, The charge for each PVBC (EPSQ and IPSQ), calculated in 2 ms bins by integrating the current in a sliding 100 ms window centered around that bin. Summary data in ***E.1*** and ***E.4*** thus represent these curves at the y-intercept. Asterisks indicate regions that survived Šidák’s multiple comparisons * < 0.05, ** < 0.01, *** < 0.01. ***H***, the ratios of curves in ***G*** yield the synaptic E/I ratio during SWRs. Summary data in ***F*** represents these curves at the y-intercept. ***I***, Linear regression of spike rate during SWRs vs the E/I ratio. For all plots, individual data points represent a cell. Closed circles represent cells from males, open circles females. Bar plots represent normal data with Mean ± SEM. p-values indicated above brackets.

During SWRs, PVBCs from 5xFAD mice saw a 50.1 ± 10.7% reduction in the swrEPSQ, the total excitatory synaptic charge in a 100 ms window centered around the SWR peak (t_(26)_ = 3.20, p = 0.0036, unpaired t-test; Fig. 8E.1), whereas there was no change for PVBSCs (t_(8)_ = 0.997, p = 0.348; unpaired t-test; Fig. 8E.2) or PVAACs (t_(9)_ = 1.63, p = 0.139; unpaired t-test; Fig. 8D.3). In a subset of cells (n_PVBC_ = 8 CT, 8 5xFAD), we also recorded IPSCs at 0 mV (Fig. 8A.4 for PVBCs, IPSCs were not recorded in every cell, thus there were insufficient numbers of PVBSCs and PVAACs for statistical analysis). Spontaneous inhibitory input to PVBCs was unchanged, including sIPSC frequency (t_(14)_ = 0.448, p = 0.661; unpaired t-test), amplitude (t_(14)_ = 0.339, p = 0.740; unpaired t-test; Fig. 8B.4), and kinetics (Rise: t_(14)_ = 0.263, p = 0.796; Decay: t_(14)_ = 0.672, p = 0.513; Fig. 8C.4). The total inhibitory charge during SWRs, swrIPSQ, was also no different between genotype (t_(14)_ = 0.0083, p = 0.994; unpaired t-test; Fig. E.4). This selective decrease in excitation to PVBCs during SWRs, in contrast to the observation in PCs (Fig. 5J), resulted in a significant decrease in the synaptic E/I ratio during SWRs (t_(14)_ = 2.70, p = 0.017, unpaired t-test), though not during spontaneous periods (t_(14)_ = 0.929, p = 0.368, unpaired t-test; Fig. 8F). As with PCs, we examined the time course of synaptic charge during SWRs in PVBCs, and found a significant effect of genotype on the excitatory charge (F_(1,2500)_ = 664, p < 10^−15^; 2-way ANOVA), with bins from −32 to +42 ms relative to the SWR peak surviving multiple comparisons (Fig. 8G). There was no effect of genotype on inhibitory charge (F_(1,1400)_ = 2.12, p = 0.145, 2-way ANOVA; Fig. 8G). This resulted in a significant effect of genotype on the E/I ratio (F_(1,1300)_ = 339, p < 10^−15^, 2-way ANOVA), with an early peak that survived multiple comparisons in control above 5xFAD cells from −66 to −58 ms relative to the SWR peak (Fig. 8H), precisely the opposite effect observed in PCs (Fig. 5W-X)

Since we recorded the cell-attached spiking activity from the same PVBCs, we were next interested if reduced SWR spike rate (Fig. 7B.1) was correlated with altered synaptic E/I ratio during SWRs. We observed a moderate positive correlation for control PVBCs (n_PVBC_ = 8, R^2^ = 0.625, F_(1,6)_ = 9.98, p = 0.020, Linear Regression), but no correlation between spike rate and E/I ratio for 5xFAD PVBCs (n_PVBC_ = 8, R^2^ = 0.0092, F_(1,6)_ = 0.056, p = 0.822; Fig. 8I), suggesting that not only is excitatory synaptic input during SWRs reduced, potentially through altered PC-PVBC connectivity, but there are also deficits in PVBC input-output function, consistent with prior studies of intrinsic PV cell dysfunction (Verret et al., 2012).

## Discussion

Here we identified a selective reduction in PVBC activity in a model of AD, while PVBSCs, PVAACs, and excitatory PCs were relatively spared. By investigating the synaptic input and spike output of these cell types, we present a careful description of hippocampal micro-circuitry alterations in early amyloid pathology (Fig. 9) during activity critical for memory consolidation (i.e. SWRs). PVBCs displayed a reduced synaptic E/I ratio during SWRs (Fig. 8F), driven by a reduction in excitatory synaptic input (Fig. 8E.1). PVBCs spiked less during SWRs (Fig. 7B.1), thus providing reduced inhibitory control to excitatory PCs. In contrast, PCs displayed an increased synaptic E/I ratio (Fig. 5S-T), with differences between superficial and deep PCs. sPCs saw a greater reduction in inhibitory input than dPCs (Fig. 5Q-R), and also displayed an increased probability of spiking during SWRs (Fig. 4E-F). As the strong inhibition PVBCs provide is critical for the selection of PC ensembles (Klausberger and Somogyi, 2008; Ellender et al., 2010), this selective reduction may explain the enlarged PC ensembles (Fig. 3E), aberrant cellular participation (Fig. 3L-M), and more frequent and larger amplitude SWRs in 5xFAD mice (Fig. 2C-D). Intriguingly, the increase in SWRs appears detrimental, despite their role in memory consolidation, as we also observed spatial memory deficits (Fig. 1C). Considering the surprising role SWRs play in down-regulating synapses and reducing memory-irrelevant activity (Norimoto et al., 2018), these aberrant SWRs may be interfering with memory-relevant reactivations. Several mechanisms likely contribute to the increase in SWRs. The reduction of sIPSCs that sPCs receive between SWRs (Fig. 5K) may permit the buildup of excitation necessary for SWR initiation. The decreased duration of SWRs (Fig. 2D) may more quickly reset the system for subsequent events. A more complete description would also require investigation of CA3 and CA2 micro-circuitry.

**Figure 9:**
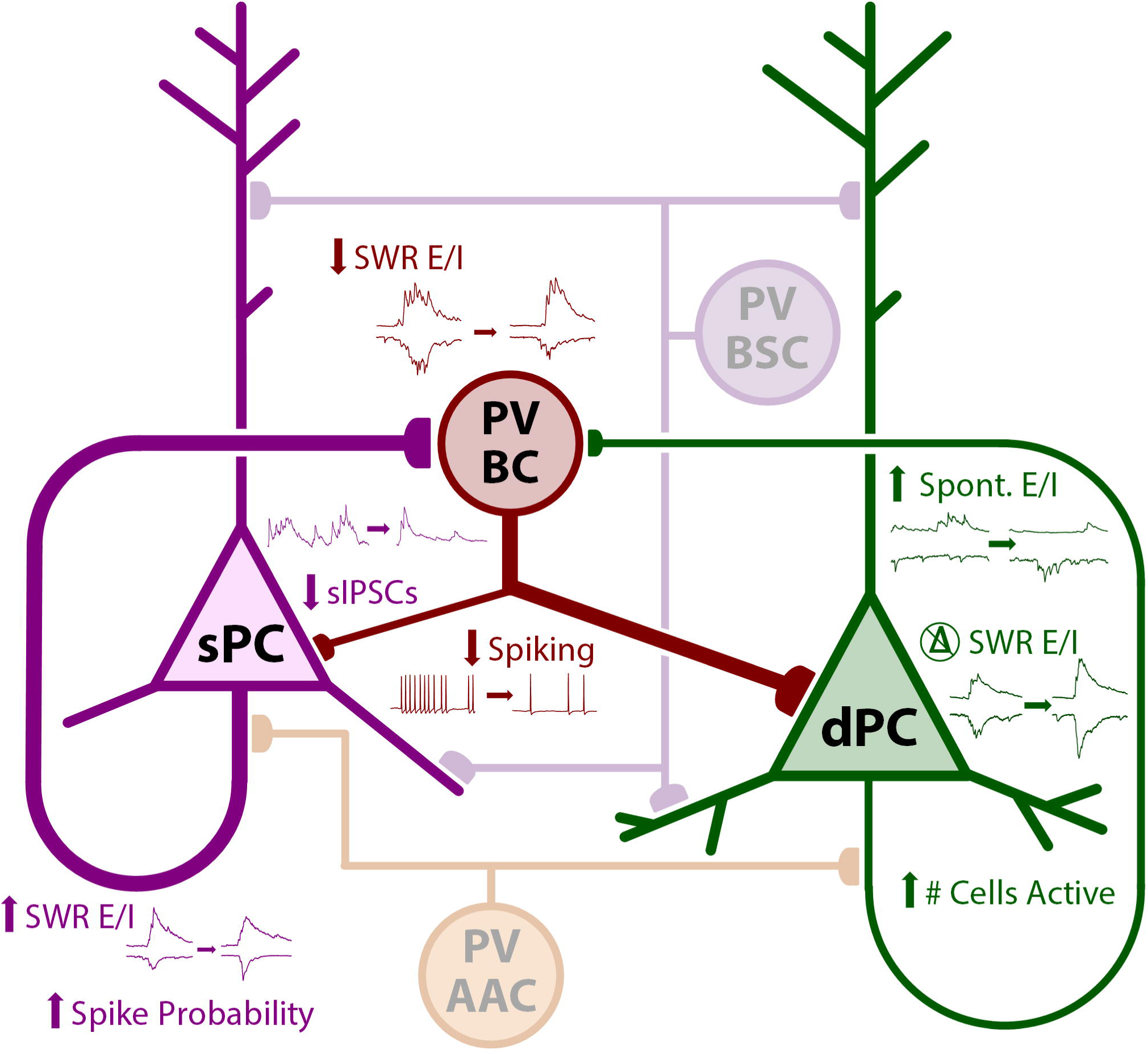
Schematic of alterations to the CA1 micro-circuit in 5xFAD mice. In PVBCs, the synaptic E/I ratio during SWRs was reduced, with a reduction in spike rate during SWRs. In sPCs, there was a reduction in sIPSCs, an increased E/I ratio during SWRs, and an increased probability of spiking during SWRs. In dPCs, there was an increase in synaptic E/I during spontaneous periods (between SWRs), no change in the E/I ratio during SWRs (with concomitant increases in excitation and inhibition), and enlarged ensembles.

### Hyperactivity and inhibitory disruption in Alzheimer’s disease

Our findings are consistent with growing evidence linking hyperactivity and Aβ aggregation (Zott et al., 2018). Hippocampal hyperactivity is seen in mouse models of early amyloid pathology as increased seizure risk (Palop et al., 2007) and enlarged ensemble activity (Busche et al., 2012). Our results extend these findings to the study of SWRs. In addition, several studies have demonstrated that a preferential disruption to inhibitory cells underlies hyperactivity (Verret et al., 2012; Hazra et al., 2013; Mahar et al., 2016; Hijazi et al., 2019). In the clinical population, seizures are more prevalent and associated with earlier onset of cognitive decline in amnestic MCI patients (Vossel et al., 2013). Additionally, task-related engagement of the hippocampus, as tested via event-related fMRI, indicates hippocampal hyper-activation in MCI patients relative to aged controls, while more progressed AD patients experience hypo-activation (Dickerson et al., 2005; Pariente et al., 2005). These studies suggest our observed phenotype may better model mild cognitive impairment rather than fully progressed AD.

### Sharp wave ripple alterations in aging and disease

In normal aging, SWR event and ripple frequency decrease (Wiegand et al., 2016; Cowen et al., 2018), contrasting with the phenotype we observed in younger mice. While it is accepted that AD is distinct from accelerated aging (Toepper, 2017), similar decreases in SWRs have been observed in aged AD models. In an apoE-ε4 knockin model of sporadic AD, SWR event frequency and gamma power are reduced in 12-18 mo mice (Gillespie et al., 2016; Jones et al., 2019). Similarly, in 9-12 mo TgF344-AD rats, SWR event frequency, power, and gamma power are reduced (Stoiljkovic et al., 2018). In younger adults the findings are somewhat mixed. As seen in a cohort of six 3 mo 5xFAD mice, SWR event frequency and gamma power during non-theta periods are reduced (Iaccarino et al., 2016), the opposite phenotype we observe in slice. In 2-4 mo rTg4510 mice, SWR event frequency is unchanged while amplitude and ripple power are reduced (Ciupek et al., 2015). Another study has suggested a failure of PC-PV circuits in amyloid pathology is due to regulation of the GluA4 AMPA receptor, resulting in more frequent SWRs with reduced ripple frequency in 3-4 mo APP_swe_/PS1ΔE9 mice (Xiao et al., 2017). Propagation of SWRs also appears disrupted; 3 mo APP-PS1 mice show impaired propagation from CA3 to CA1 correlated with increased immunoreactivity for PV (Hollnagel et al., 2019).

One notable difference between our study and others is the acute slice versus *in vivo* preparation. Slice electrophysiology permits a careful study of the synaptic inputs to different neuronal sub-types during SWRs, an infeasible task *in vivo*. It is unknown if awake versus asleep SWR are differentially affected in AD, and it is unclear which, if either, are better modeled in slice. The choice of AD model also surely has implications. The 5xFAD model has several advantages over other mouse models in replicating human disease: a high ratio of Aβ_42_ over Aβ_40_, memory impairment, and neurodegeneration in later disease progression. However, it also has limitations. As with most AD models, it only models the familial variant of the disease, while the sporadic variant accounts for most human cases. Moreover, the presence of five APP/PS1 gene mutations would certainly never be observed in a patient. Another disadvantage is APP overexpression, common to all first-generation transgenic models. Some have noted electrophysiological alterations are more related to APP overexpression than Aβ aggregation (Born et al., 2014). We attempted to address this by examining a 1 mo cohort (preceding plaque aggregation), in which APP overexpression would still presumably have an effect, a cohort in which we observed no alterations (Fig. 1C,2C). Fewer studies exist of second-generation APP knockin models which address the APP overexpression problem, yet have less pronounced disease phenotypes (Sasaguri et al., 2017). In one relevant study, sIPSC amplitude from putative PV cells is decreased in parietal cortex PCs from 18-20 mo App^NL-F^ mice, although synaptic E/I balance is unchanged (Chen et al., 2018). It will be critical to study hippocampal network alterations, both at the level of the micro-circuit and behaving animal, in novel AD mouse models with greater validity.

### Potential mechanisms for PVBC deficit

Several mechanisms may underlie PVBC dysfunction in amyloid pathology. Intrinsic factors including downregulation of the voltage-gated Na_v1.1_ channel can explain decreased PV cell excitability and cortical network hyperactivity (Verret et al., 2012). It remains to be shown if different PV sub-types are differentially affected, but according to the Allen Brain Institute Cell Types Database, all identified murine PV clusters express the Scn1a gene encoding Na_v1.1_ (Lein et al., 2007). Another potential mechanism is loss of peri-neuronal nets (PNNs), part of the extracellular matrix surrounding soma and proximal dendrites that preferentially ensheathe PV cells (Kwok et al., 2011; Sorg et al., 2016). The function of PNN is incompletely understood, but both enhances PV cell excitability and activity (Slaker et al., 2015; Balmer, 2016). In a prior study, we observed degradation of PNN increases SWR event frequency by ∼50% (Sun et al., 2017), a similar magnitude effect as the current study. Additionally, the selective reduction of excitatory input to PV cells (Fig. 8E.1) is consistent with the decrease in miniature EPSCs in PV cells in a brevican knockout, a major component of PNN (Favuzzi et al., 2017). Moreover, PNN staining is reduced in 3 mo Tg2576 mice (Cattaud et al., 2018). PNNs are more selectively located around PVBCs (>90%) than PVBSCs (25-50%) or PVAACs (<10%) (Yamada and Jinno, 2015), potentially explaining the specificity we observe. However, our experiments would also be consistent with upstream deficits in excitatory input to PV cells. In 4 mo Tg2576 mice, there is a preferential degeneration of direct entorhinal input to CA1 PV cells; optogenetic restoration of this input rescues synaptic and spatial learning deficits (Yang et al., 2016). The overall cause of disruption to PV cells is likely from several contributing factors, and further studies are required to identify the most salient impairment, and thus the most promising avenue for intervention.

### Implications of PVBC specific deficit

Here, we identified a preferential impairment in PVBC function, concurrent with altered network activity. The selectivity of synaptic alterations in PVBCs suggest they may be a promising target for intervention to restore hippocampal network activity in early amyloid pathology. Given the rapid evolution of tools to manipulate activity in a cell-type specific manner, this finding is of particular importance considering optogenetic (Iaccarino et al., 2016) and chemogenetic (Hijazi et al., 2019) approaches to ameliorate memory decline in AD. A limitation of these strategies is that the PV^Cre^ driver will also target PVBSCs and PVAACs, potentially leading to unintended off-target consequences in temporal sequencing. Novel combinatorial approaches utilizing both Cre and Flp (He et al., 2016) provide a promising avenue to selectively manipulate distinct neuronal sub-types. For example, the Nkx2.1^CreER^;LSL^Flp^ mouse would provide an efficient means to record PVAAC activity, in which we saw small alterations that did not reach significance. In conclusion, this study investigated the synaptic input and spiking output of distinct PC and PV cell-types within CA1 micro-circuitry over the course of SWR events, providing insight into synaptic deficits in early amyloid pathology, and informing future attempts to manipulate the hippocampal micro-circuit.

## Acknowledgements

AC was supported by NIH/NIA F31 AG062030, NIH/NCATS TL1 TR001431, and the ARCS Foundation. LB was supported by NIH/NINDS T32 NS041218. DP and SV were partially supported by NIH/NIA RF1 AG056603. KC and SV were partially supported by NIH/NINDS R01 NS108810. The content is solely the responsibility of the authors and does not necessarily represent the official views of the National Institutes of Health.

